# Single-cell profiling of Ebola virus infection *in vivo* reveals viral and host transcriptional dynamics

**DOI:** 10.1101/2020.06.12.148957

**Authors:** Dylan Kotliar, Aaron E. Lin, James Logue, Travis K. Hughes, Nadine M. Khoury, Siddharth S. Raju, Marc H. Wadsworth, Han Chen, Jonathan R. Kurtz, Bonnie Dighero-Kemp, Zach B. Bjornson, Nilanjan Mukherjee, Brian A. Sellers, Nancy Tran, Matthew R. Bauer, Gordon C. Adams, Ricky Adams, John L. Rinn, Marta Melé, Garry P. Nolan, Kayla G. Barnes, Lisa E. Hensley, David R. McIlwain, Alex K. Shalek, Pardis C. Sabeti, Richard S. Bennett

## Abstract

Ebola virus (EBOV) causes epidemics with high case fatality rates, yet remains understudied due to the challenge of experimentation in high-containment and outbreak settings. To better understand EBOV infection *in vivo*, we used single-cell transcriptomics and CyTOF-based single-cell protein quantification to characterize peripheral immune cell activity during EBOV infection in rhesus monkeys. We obtained 100,000 transcriptomes and 15,000,000 protein profiles, providing insight into pathogenesis. We find that immature, proliferative monocyte-lineage cells with reduced antigen presentation capacity replace conventional circulating monocyte subsets within days of infection, while lymphocytes upregulate apoptosis genes and decline in abundance. By quantifying viral RNA abundance in individual cells, we identify molecular determinants of tropism and examine temporal dynamics in viral and host gene expression. Within infected cells, we observe that EBOV down-regulates *STAT1* mRNA and interferon signaling, and up-regulates putative pro-viral genes (e.g., *DYNLL1* and *HSPA5*), nominating cellular pathways the virus manipulates for its replication. Overall, this study sheds light on EBOV tropism, replication dynamics, and elicited immune response, and provides a framework for characterizing interactions between hosts and emerging viruses in a maximum containment setting.

## Introduction

Ebola virus (EBOV) is among the world’s most lethal pathogens, with an estimated case fatality rate of 66% in the recent epidemic in the Democratic Republic of the Congo (Ilunga Kalenga et al., 2019; World Health Organization, 2019) and 40% in the 2013–2016 epidemic in West Africa (Lo et al., 2017). EBOV infection in humans causes Ebola virus disease (EVD), characterized by fever, malaise, muscle aches, and gastrointestinal distress, rapidly progressing to coagulopathy, shock, and multi-organ failure (Malvy et al., 2019). While recently developed vaccines (Kennedy et al., 2017) and monoclonal antibody therapeutics (Mulangu et al., 2019) have shown great promise for preventing and treating EVD, case fatality rates in treated patients still exceed 30%, highlighting the need for further research into disease pathogenesis.

Studies of EVD pathogenesis, while paramount, face numerous logistical challenges which have limited their scope relative to studies of other pathogens. Experiments involving live EBOV require maximum containment (e.g., biosafety level 4 [BSL-4]) and therefore are restricted to a small number of highly specialized research facilities. *In vivo* studies are especially challenging: human EVD is difficult to study in the midst of deadly outbreaks in resource-limited settings, necessitating animal models. However, commonly used laboratory mouse lines are resistant to naturally occurring EBOV isolates, limiting their utility for research (Bray, 2001; Rasmussen et al., 2014). Moreover, rodents and other small animal models such as ferrets lack the primate-specific NPC1 genotype (Diehl et al., 2016), the key cellular receptor for EBOV entry (Carette et al., 2011; Côté et al., 2011), and do not always recapitulate human EVD-like disease (Bray et al., 2001; Geisbert et al., 2002). EVD in nonhuman primates (NHPs) most closely resembles human EVD (Bennett et al., 2017; Geisbert et al., 2015; St Claire et al., 2017), but NHP studies are often limited to small sample sizes, which reduces power to identify statistically significant trends and to discern meaningful inter-individual variability.

The two predominant approaches to studying EVD – analyzing infected cells in culture and infected animals *in vivo* – have revealed important, if somewhat contradictory, aspects of how EBOV impacts the host immune system. In cell culture, EBOV infects myeloid cells, potently inhibiting both production of type 1 interferon (Basler et al., 2003; Gupta et al., 2001; Harcourt et al., 1998) and signal transduction downstream of interferon receptors (Harcourt et al., 1999; Kash et al., 2006; Leung et al., 2006; Reid et al., 2008). Under-activation of this key innate antiviral response hinders the ability of antigen-presenting cells to activate the adaptive immune system to combat infection (Bosio et al., 2003; Lubaki et al., 2013; Mahanty et al., 2003), a key determinant of fatal outcomes (Baize et al., 1999) and could be due to reduced presentation of viral proteins by antigen-presenting cells (Lüdtke et al., 2016). In contrast to these culture-based findings, EVD *in vivo* is characterized by high fever and dramatic up-regulation of hundreds of interferon stimulated genes (Caballero et al., 2016; Liu et al., 2017; Reynard et al., 2019; Speranza et al., 2018), correlating with the release of dozens of inflammatory cytokines (Caballero et al., 2016; Reynard et al., 2019; Wauquier et al., 2010), suggesting that an aberrant over-activation of innate and adaptive immunity underlies much of EVD pathology, rather than solely virus-mediated cytotoxicity (Geisbert et al., 2003a, 2003b).

High-throughput single-cell technologies, such as single-cell RNA-sequencing (scRNA-Seq) and protein quantification by CyTOF (Bendall et al., 2011), have made it possible to analyze the response of individual cells to viral infection at unprecedented resolution (Hamlin et al., 2017; Hein and Weissman, 2019; Newell et al., 2012; O’Neal et al., 2019; Russell et al., 2018, 2019; Steuerman et al., 2018; Zanini et al., 2018a; Zhao et al., 2020). By generating mRNA or protein profiles for thousands of cells in a sample, these methods can quantify the cell-type composition and expression programs of individual cells -- signals that are obscured in bulk measurements. By quantifying viral RNA within individual cells, scRNA-Seq allows comparison of gene expression between infected and uninfected cells in a diseased host (*i.e.*, bystander cells), which can yield a far more nuanced view of host (Steuerman et al., 2018) and viral (Hein and Weissman, 2019) gene expression within infected cells. Further, this approach can be used to disentangle the direct effects of EBOV infection within a cell from the effect of the inflammatory cytokine milieu. However, many scRNA-Seq technologies require droplet generator devices and inactivation protocols that can be challenging to establish in a maximum containment facility. As a result, such approaches have yet to be applied to a risk group 4 (RG-4) pathogen such as EBOV. Furthermore, high-volume exhaust, superheated components, and other aspects of CyTOF instrumentation make these devices incompatible with installation in maximum containment facilities (Logue et al., 2019), necessitating development of new protocols compatible with sample inactivation to study RG-4 pathogens.

Here, we describe the first investigation of an RG-4 agent under maximum containment with high-dimensional single-cell technologies. We apply CyTOF and Seq-Well--a portable single-cell RNA-seq platform (Gierahn et al., 2017; Hughes et al., 2019)--to a combined total of 90 peripheral blood mononuclear cell (PBMC) samples (90 by CyTOF, 28 by Seq-Well) collected from 21 rhesus monkeys prior to infection or at multiple timepoints following lethal EBOV challenge *in vivo*. We further inoculated PBMCs with EBOV *ex vivo*, using pre-defined experimental parameters, and profiled their gene expression with Seq-Well. These data allow us to dissect host-virus interactions and comprehensively catalog changes in cell-type abundance and cell state over the course of EVD. Moreover, as EBOV harbors an RNA genome and transcribes polyadenylated mRNAs, we were able to detect viral RNA within individual cells, allowing us to define EBOV tropism with high resolution and identify EBOV-associated transcriptional changes in putative pro- and antiviral genes.

We find that EVD leads to widespread changes in the circulating monocyte populations *in vivo*, both in NHPs as well as in acute human infections, with replacement of conventional monocyte subsets with a highly proliferative monocyte precursor population and a macrophage-like population that is enriched for EBOV-infected cells. Furthermore, by comparing infected and uninfected bystander monocytes, we resolve the apparent contradiction between *in vivo* and *in vitro* studies of EBOV. We find that bystander cells of all major immune cell types express an interferon response program, but that this response is suppressed specifically within infected monocytes *in vivo*, consistent with previous studies in culture. In addition to down-regulating host antiviral genes, we observe that EBOV drives the up-regulation of candidate pro-viral genes, such as *DYNLL1* and *HSPA5*, within infected cells. Taken together, this dataset constitutes a unique resource for the study of EBOV, enabling the study of host immune response in infected and bystander cells across cell types, in natural (*in vivo*) and experimentally controlled (*ex vivo*) contexts.

## Results

### Single-cell characterization of RNA and protein expression in circulating immune cells from Ebola virus infected rhesus monkeys

To comprehensively profile EBOV-induced immune dysfunction *in vivo*, we collected peripheral immune cells from rhesus monkeys prior to infection and at multiple days post-infection (DPI), corresponding to several stages of acute EVD (Figure 1). Cohorts of ≥3 nonhuman primates (NHPs) were sacrificed as baseline uninfected controls (B), at pre-defined DPI, or upon reaching pre-determined humane euthanasia criteria. These cohorts were recently characterized for viral load, clinical score, blood chemistry (Bennett et al. in submission) and liver pathology (Greenberg et al., 2020). Viral load first became detectable in all animals on DPI 3, preceding detectable clinical signs (*e.g.*, fever) by 1–2 days (Figure 2A). Clinical signs of EVD progressed until humane euthanasia criteria were uniformly reached between DPI 6–8 (Figure S1A). For each NHP, cells collected multiple times throughout disease were used for CyTOF, while cells collected prior to infection and at sacrifice were used for CyOF and Seq-Well (Figure S1B).

**Figure 1.**
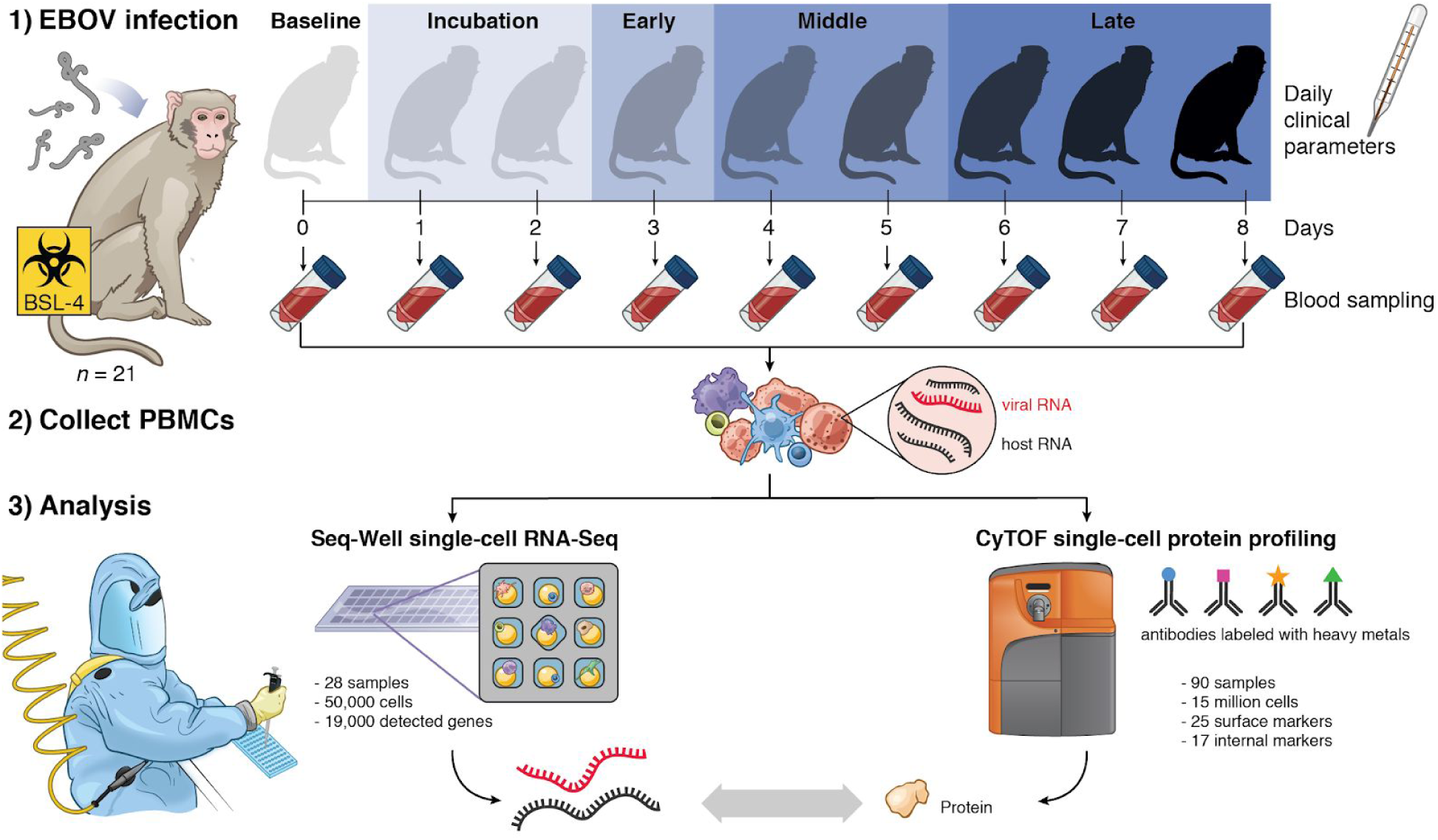
Studying lethal Ebola virus disease in rhesus monkeys using single-cell profiling of RNA and protein expression in peripheral immune cells. Serial sampling study design for investigating the host response to Ebola virus (EBOV) infection in rhesus monkeys. Under biosafety level 4 (BSL-4) containment, we collected blood samples from a total of 21 animals at multiple days post inoculation, extracted peripheral blood mononuclear cells (PBMCs), and profiled single-cell transcriptomes and 42 protein markers using Seq-Well and CyTOF. Seq-Well quantifies both host (black) and viral (red) RNA expression, allowing comparisons between infected and bystander cells. We also assessed daily clinical parameters for each animal, including body temperature, clinical signs, and body weight, and obtained complete blood counts for each blood draw. See also Figure S1 and **Table S1** for a complete listing of CyTOF and Seq-Well samples.

**Figure 2.**
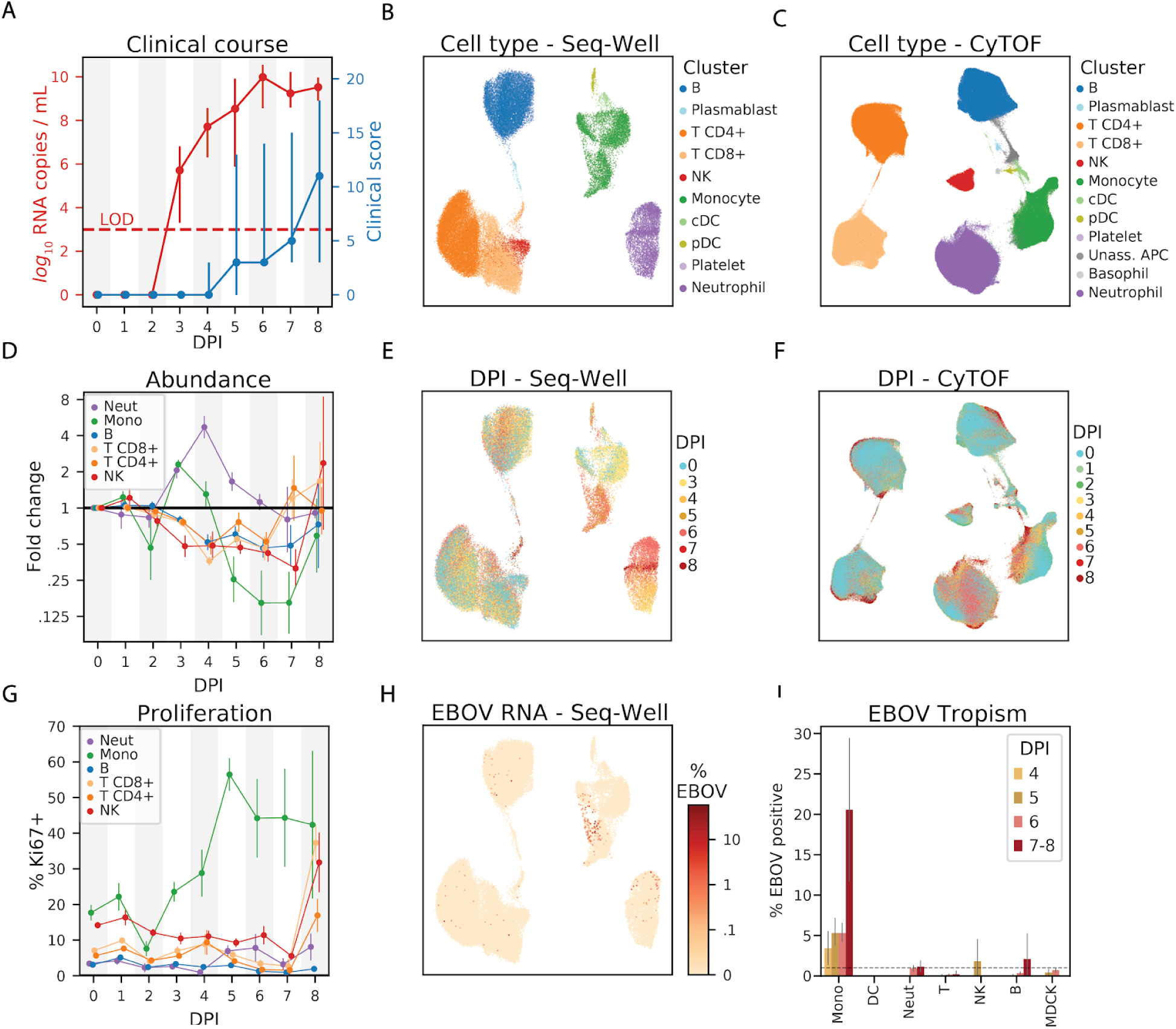
Single-cell profiling reveals changing cell-type abundance, proliferation rate, and infection status throughout EVD. (**A**) Time course of viral load (red, left *y*-axis, log_10_ scale) and clinical score (blue, right *y*-axis). Markers denote the mean and error bars denote the minimum and maximum values. LOD, limit of detection of viral load by RT-qPCR assay. See also Figure S1. (**B and C**) Uniform Manifold Approximation and Projection (UMAP) embedding of Seq-Well (B) and CyTOF (C) data, colored by annotated cluster assignment. See also Figures S2 and S4. (**D**) Fold change (log_2_ scale) in the absolute abundance (cells / µL of whole blood) of each cell type relative to baseline based on clustering of CyTOF data. Error bars denote the mean ± 1 standard error. See also Figures S5A and S5B. (**E and F**) UMAP embedding of Seq-Well (E) and CyTOF (F) data, colored by the day post infection (DPI) on which each cell was sampled. Percentage of Ki67-positive cells (CyTOF intensity > 1.8, arbitrary units) of each cell type. Error bars denote the mean ± 1 standard error. See also Figures S5C and S5D. (**G**) UMAP embedding of Seq-Well data, colored by the percentage of cellular transcripts mapping to EBOV. (**H**) Percentage of infected cells by cell type based on Seq-Well expression profiles. Dashed line denotes the 1% false positive rate threshold used for calling infected cells. See also Figure S2G.

After standard quality control filters (**Materials and Methods**), we obtained single-cell transcriptomes and 42-protein CyTOF profiles from ∼58,000 and ∼15,000,000 PBMCs, respectively. We visualized these data with uniform manifold approximation and projection (UMAP) non-linear dimensionality reduction (Becht et al., 2018; McInnes et al., 2018) (Figures 2B, 2C, 2E, 2F, **and** 2H). Unsupervised clustering of either the transcriptomes or a down-sampled set of 1,100,000 protein profiles (**Materials and Methods**) yielded clusters that could be readily identified as the major circulating immune cell types using well-known RNA and protein markers (Figures 2B, 2C, S2A, **and** S2B). The CyTOF-based clustering gave concordant results with an approach based on manual gating of conventional cell-type marker genes (Figures S3A and S3B**, Materials and Methods**). After batch correction of the CyTOF data and integration of the transcriptomes to adjust for technical sources of variation **(Materials and Methods**), samples were well-distributed across cell-type clusters (Figures S2C–S2F) but separated by DPI (Figures 2E **and** 2F), suggestive of dynamic cell states over the course of disease progression. By sub-clustering within broad cell-type categories, we further identified cell subtypes based on the expression of identifying marker genes (Figure S4).

### Cell-type abundance, proliferation, and EBOV infection rates vary throughout EVD

In addition to the major PBMC cell types, a cluster of immature neutrophils emerged during EVD, marked by high gene expression of *CD177* and *SOD2*, and protein expression of CD66 and CD11b. Though neutrophils are typically removed during density-based PBMC isolations, immature neutrophils (*i.e.*, band cells) – which are less dense than mature polymorphonuclear neutrophils – can be released from the bone marrow and co-isolate with PBMCs in infectious and autoimmune inflammatory conditions (Carmona-Rivera and Kaplan, 2013; Darcy et al., 2014; Deng et al., 2016), including during EVD (Eisfeld et al., 2017). Neutrophils were almost entirely absent from baseline samples in our data but constituted a high proportion of cells in late EVD samples (scRNA-Seq: 0.2% of baseline cells compared to 65.1% of late EVD cells; CyTOF: 9.3% of baseline compared to 49.8% of late EVD; Figures S5A **and** S5B), which supports the hypothesis that band cells are released into the periphery from the bone marrow in response to cytokines elicited during EVD.

Next, we quantified absolute abundance of each cell type over the course of EVD by combining CyTOF data with complete blood counts (CBC) (**Materials and Methods**) (Bennett et al. in submission). CBC provided direct neutrophil, lymphocyte, and monocyte abundances, and we integrated CBC and CyTOF data to obtain differential abundances of the lymphocyte cell types. Cell-type percentage estimates based on CyTOF were in general agreement with those based on scRNA-Seq (Figure S5A).

In agreement with previous NHP studies (Ebihara et al., 2011; Fisher-Hoch et al., 1985), there was a >5-fold increase in neutrophil abundance by DPI 4 relative to baseline, before levels returned to baseline in late EVD (*p* < .05 for DPI 3–4, *p* = 0.059 for DPI 5, Wilcoxon signed-rank test, Figures 2D **and** S5B). Also consistent with previous studies, we observed a marked decrease in lymphocyte abundance with NK cells declining one day before the other cell types (*p* < 0.05 on DPI 3–6 for B, NK, CD8+ T, and CD4+ T; except for CD4+ T on DPI 5, Wilcoxon signed-rank test). Interestingly, all lymphocyte populations slowly recovered after DPI 4 (Figure S5B). Monocyte abundance initially increased >2-fold before declining precipitously between DPI 4 and 5.

Changes in circulating cell-type abundance could reflect cell proliferation and/or death, as well as movement of cells into and out of bone marrow, lymph, and tissues. While we were unable to directly quantify rates of death or movement between different compartments, we estimated the fraction of actively dividing cells using the proliferation marker Ki67 (encoded by the gene *MKI67*) in both the CyTOF and scRNA-Seq data, and found good agreement between the two modalities (Figure S5C).

The fraction of Ki67+ monocytes increased dramatically from 17% at baseline to 56% at DPI 5 and remained >40% for the remainder of disease (*p* = 1.1×10^-5^ rank-sum test of DPI 5-8 vs. baseline samples), suggesting an increase in proliferation (Figures 2G **and** S5D). By contrast, neutrophil proliferation remained roughly constant (Figures 2G **and** S5D) despite the dramatic changes in abundance (Figures 2D **and** S5B), further evidence that immature neutrophils were released from the bone marrow during disease (Summers et al., 2010). Intriguingly, the fraction of dividing T and NK cells stayed relatively constant for most of the time course but increased dramatically on DPI 8 for both of the NHPs (out of 6 total) that survived until then, both by RNA (Figure S5C) and protein levels (*p* = 0.022 rank-sum test of DPI 8 vs. baseline for NK, CD8+ T, and CD4+ T, Figure S5D). Proliferation is a core component of effective T-cell mediated viral clearance, but requires time for activated T cells to accumulate; the observation that significant proliferation of circulating T cells only occurred in the 2 animals that survived until the latest DPI suggests that those animals may have begun to mount a T-cell response.

Not all cell types support EBOV entry and replication; here, we were able to identify which cells were infected *in vivo* using scRNA-Seq because EBOV has an RNA genome and produces poly-adenylated mRNA transcripts (Figure 2H). However, uninfected cells may also contain EBOV reads due to ambient RNAs that contaminate single-cell profiles (Fleming et al., 2019; Young and Behjati, 2018). We therefore developed a statistical approach to identify infected cells as those that contain more EBOV transcripts than would be expected by chance, based on the relative abundance of EBOV-mapped transcripts in a cell and the amount of ambient RNA contamination (**Materials and Methods**). This allowed us to control the false positive rate (FPR) at a pre-specified level while maximizing power to call infected cells. At a FPR of 1%, we were well-powered to identify an infected cell when ≥1% of its transcripts mapped to EBOV, and estimated an average sensitivity of 51% when ≥0.1% of cellular transcripts derived from EBOV (Figure S2G) though the sensitivity for a given cell depends on read depth and other parameters (**Materials and Methods**). In addition, we spiked uninfected Madin-Darby canine kidney (MDCK) cells into a subset of PBMC samples to serve as a negative control (**Table S1, Materials and Methods**).

Monocytes comprised the main infected cell population *in vivo*, first detectable at DPI 4, with an increasing fraction of infected monocytes thereafter (Figure 2I). Consistent with previous studies, T cells, B cells, and neutrophils were not identified as infected more often than would be expected by chance (1% FPR threshold), nor more often than MDCK control cells. We did not observe any infected plasmacytoid (pDC) or conventional dendritic cells (cDC) in circulation, though infected DCs have been observed in culture and in lymph nodes (Geisbert et al., 2003c) (see **Discussion**).

### Interferon response drives gene expression programs across multiple cell types

Having examined temporal shifts in the frequency of each immune cell type, we next sought to comprehensively catalog changes in their respective gene expression profiles throughout EVD. To increase statistical power to detect differentially expressed genes, we grouped cells into EVD stages based on clinically relevant phenomena: “incubation” which precedes detectable viral load or clinical signs (DPI 1 and 2; CyTOF only), “early” when there is detectable viral load but no clinical signs (DPI 3), “middle” when there is detectable viral load and clinical signs (DPI 4 and 5), and “late” when animals uniformly reached human euthanasia criteria (DPI 6–8) (Figure 1).

We compared transcriptomes of cells from each EVD stage to baseline for each cell type individually (**Materials and Methods**). This identified 1,437 differentially expressed genes with an FDR corrected *q*-value < 0.05 and a fold-change of greater than 30% in at least one cell type and stage (**Table S2**). To identify patterns of gene expression associated with cell type and time, we performed unsupervised clustering of the differential expression signatures and identified 11 modules of genes sharing similar patterns of changes (Figures 3A **and** 3B, **Table S3**, **Materials and Methods**). We excluded neutrophils, pDCs, cDCs, and plasmablasts because of small sample sizes. Three modules which we term “Global” were broadly up or down-regulated across cell types, and the remaining modules were cell-type specific.

**Figure 3.**
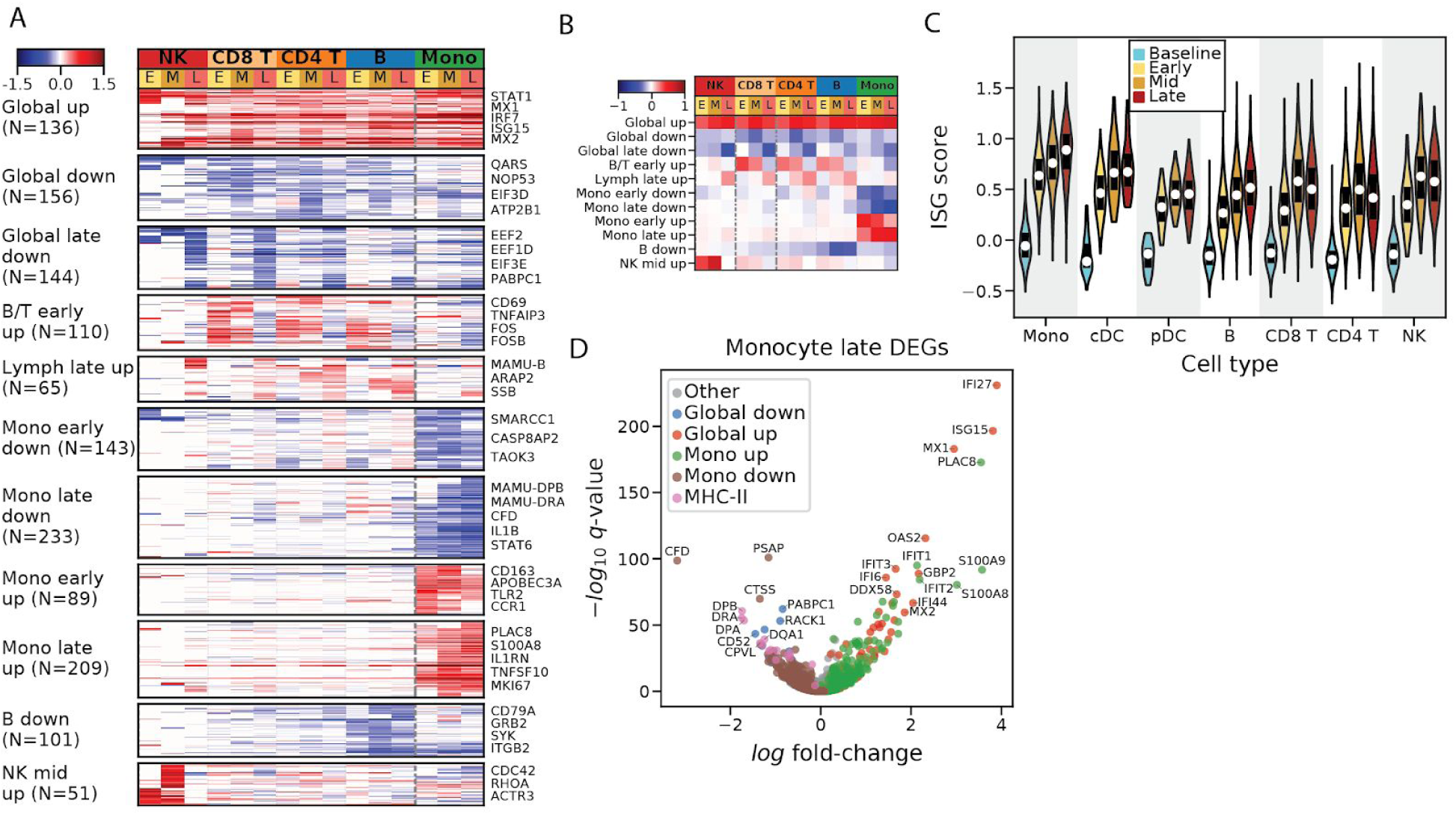
Patterns of differential expression across disease stages and cell types in EVD. (**A**) Fold changes (natural log scale) of 1,430 differentially expressed genes (rows) in each cell type at early (E), middle (M), and late (L) stages of disease, relative to baseline levels, with insignificant values (*p* > 0.2) set to 0. Genes were grouped into modules through unsupervised *k*-means clustering. See also **Tables S2 and S3**. (**B**) Same as A but displaying the average log fold-change of each module. (**C**) Distribution of interferon stimulated gene (ISG) scores for each cell type at baseline (blue), early (yellow), mid (orange), or late (red) EVD. White markers denote the median and bars denote the interquartile range. See also Figures S6A and S6B. (**D**) Volcano plot of differential expression between monocytes in late EVD compared to baseline. Genes are colored by membership in the “Global down” or “Global late down” modules (blue), “Global up” module (red), “Mono early up” or “Mono late up” modules (green), “Mono early down” or “Mono late down” modules (brown), or if they are annotated as an MHC class II gene (purple).

The “Global up” module contained 136 genes, consisting mostly of regulators and targets of the interferon (IFN) alpha (α) and gamma (γ) signal transduction cascade such as *STAT1*, *IRF7*, *MX1*, and *ISG15*. Gene sets labeled “response to interferon alpha”, and “response to interferon gamma” were significantly enriched in this module (IFNα: OR = 69.5, *q* = 8×10^-39^; IFNγ: OR = 45.9, *q* = 1×10^-39^; Fisher’s exact test; **Table S3**). The emergence of an IFN response was consistent with another observation: *IFNγ* mRNA rises >10-fold in CD8+ T-cells from an average of 0.4 transcripts per ten thousand (TP10K) at baseline to 5.0 at mid stage EVD. Concurrently, type 1 IFN (α/β) mRNAs rose from undetectable at baseline to 0.03–0.05 TP10K at late EVD in monocytes, along with a large number of other cytokines (Figure S6A). However, the increase in type 1 IFN mRNAs was not statistically significant, as IFN mRNAs are expressed transiently (Lin et al., 2011). To further characterize the dynamics of the IFN response, we assigned an interferon stimulated gene (ISG) score to each cell, reflecting the average expression of literature-annotated ISGs that overlap with the “Global up” module (**Table S3, Materials and Methods**). The median ISG score increased substantially throughout EVD across each cell type (*p* < 1×10^-5^ for all cell types and periods, rank-sum test, Figure 3C).

As there is substantial overlap between the genes stimulated by IFNα and IFNγ, we sought to determine if one cytokine predominated, or if both acted independently. We therefore identified genes that were annotated as regulated by IFNα but not IFNγ (*i.e.*, uniquely IFNα-regulated) and vice versa. Both uniquely IFNα- and uniquely IFNγ-regulated genes were significantly enriched in the “Global up” module (*q* < 0.01, **Table S3**), with a larger fold-change for the uniquely IFNα-regulated genes (IFNα OR = 20.9, IFNγ OR = 16.6). This pattern held true for each cell type and EVD stage separately (Figure S6B). These results suggest that both IFNα and IFNγ substantially and independently influenced the gene expression profiles of circulating cells during EVD.

The “Global late down” module contained 144 genes that were predominantly down-regulated across cell types during late EVD. It contained numerous regulators of translation initiation and elongation (e.g. *EEF2*, *EEF1D*, *EIF3E*, and *PABPC1*; REACTOME_TRANSLATION gene set enrichment *q* = 5.2×10^-7^, **Table S3**), which is consistent with a core antiviral function of I FN being to down-regulate translation (Li et al., 2015). The “Global down” module contained 156 genes, and similar to “Global late down”, included several other genes involved in translation (e.g. *QARS*, *NOP53*, and *EIF3D*). In addition, this module was most significantly enriched for the HALLMARK_MITOTIC_SPINDLE gene set (*q* = 8.5×10^-7^) suggesting a global down-regulation of cell cycling upon activation.

### Cell-type and temporally-specific modules underlie cell states related to clinical phenomena in EVD

After elucidating the global effects of EBOV infection on immune cells, we next investigated the transcriptional responses specific to each cell population.

The 2 modules “B/T early up” and “Lymph late up” reflect changing gene expression states of lymphocytes at different stages of acute EVD. “B/T early up” is strongly associated with the gene set HALLMARK_TNFA_SIGNALING_VIA_NFKB (*q =* 1.3×10^-9^) and is characterized by many lymphoid activation genes including the canonical marker *CD69* (Testi et al., 1994), *CD48* (McArdel et al., 2016), and the transcription factor *FOS* (Foletta et al., 1998). This module is unlikely to represent antigen-dependent activation via the BCR/TCR as it occurs in most lymphocytes and does not coincide with proliferation. Indeed, several of the top up-regulated genes, such as *GADD45B* and *ZFP36L2*, are associated with growth arrest. In addition, the 5th most enriched gene set in the “B down” module is “REACTOME_ANTIGEN_ACTIVATES_B_CELL_RECEPTOR_LEADING_TO_GENERATION_OF_SECOND_MESSENGERS” (*q* = 0.00017) suggesting a reduction in BCR activation generally. Thus, the “B/T early up” module likely represents a cytokine-mediated, non-antigen-dependent activation of lymphocytes.

The “Lymph late up” module is up-regulated in late EVD across all lymphocyte cell types. The top associated gene sets implicate DNA repair (PUJANA_ATM_PCC_NETWORK, *q* = 0.00031) and apoptosis via TRAIL (HAMAI_APOPTOSIS_VIA_TRAIL_UP, *q* = 0.00032). This latter gene set is potentially consistent with previous reports of T-cell apoptosis in EVD (Geisbert et al., 2000; Iampietro et al., 2017; Wauquier et al., 2010) and with the lymphopenia in our dataset (Figure 2D).

The “NK mid up” module is highly specific to NK cells during early EVD (Figure 3A) and is most enriched for “MARSON_BOUND_BY_FOXP3_STIMULATED” (*q = 0.0*71) and “BIOCARTA_CDC42RAC_PATHWAY” (*q* = 0.08, **Table S3**). *FOXP3* expression is characteristic of invariant NK cells (Engelmann et al., 2011) that secrete a wide variety of cytokines in response to stimulation (Krovi and Gapin, 2018). The CDC42/RAC pathway is essential for the polarization of cytolytic granules in NK cells, a requirement for effective cytotoxicity (Sinai et al., 2010; Tybulewicz and Henderson, 2009). This suggests that NK cells became activated with increased cytotoxicity in the mid EVD stage, but not in late EVD.

### Monocytes express reduced MHC class II mRNAs and proteins independent of infection status

Monocytes were of particular interest. In addition to being the preferred target of EBOV (Figure 2I), they had far more significant gene expression changes during EVD than the other cell types. 1,020 genes (11.6% of total genes tested) were differentially expressed in monocytes at one or more EVD stages versus baseline, compared to 505 genes (6.6%) for B cells, the cell type with the second most differentially expressed genes. We therefore focused our attention on characterizing monocytes in detail.

One prominent feature of the monocyte differential expression profile was the striking down-regulation of several MHC class II (MHC-II) genes by mid and late EVD (Figure 3D). Monocytes and professional antigen-presenting cells display viral antigens on MHC-II proteins at the cell surface to stimulate the adaptive immune response. While IFNγ typically up-regulates MHC-II gene and protein expression (Steimle et al., 1994), we observed decreased MHC-II on monocytes despite elevated *IFNγ* mRNA levels in T cells (Figure S6A) and widespread IFNγ transcriptional response in monocytes (Figure S6B). Previous reports have described loss of HLA-DR, one of the 4 MHC-II proteins, during EBOV infection of monocytes *ex vivo* (Hensley et al., 2002), in experimentally infected NHPs (Menicucci et al., 2017) and in human EVD cases (Lüdtke et al., 2016), similar to observations of reduced HLA-DR on monocytes in patients experiencing septic shock (Wolk et al., 2000). However, the specific MHC genes affected, the cell-type specificity, temporal dynamics, and relationship with EBOV infection status, have not been previously described.

We observed widespread changes in levels of MHC genes throughout EVD (Figure 4A). The most striking decreases occurred in MHC-II genes of monocytes (>5-fold for *DPA*, *DPB*, and *DRA* by late EVD, *q* < 1×10^-21^ for all MHC-II genes), with smaller effect-size changes in MHC-I genes (<1.7-fold increase for *A*, *A3*, and *B* at late disease, *q* < 5×10^-4^). B cells displayed modest reductions in MHC-II genes as well (>1.9-fold for *DPA*, *DPB*, *DRA*, and *DQA1* at late disease, *q* < 1×10^-22^). pDCs and cDCs showed no statistically significant reduction of any MHC-II gene (*q* > 0.05) but our dataset contained few DCs (**Table S1**), so we had less power to detect these effects. We observed a corresponding pattern in the protein levels by CyTOF: in monocytes, HLA-DR protein levels decreased to a greater extent than in the other cell types (*p* < 1×10^-61^ for monocytes in early, mid, and late stages, rank-sum test, Figure 4B), with a more modest reduction of HLA-DR in B cells at DPI 5–8 (*p* = 0.0012, rank-sum test) (Figure S6C). This phenomenon held true for each individual NHP; even as monocytes became activated, demonstrated by up-regulation of the canonical activation marker CD38 (Amici et al., 2018) (Figures 4C **and** S3D), they showed dramatic down-regulation of average HLA-DR protein expression in monocytes at DPI 5–8 versus baseline (*p* = 9.5×10^-7^, rank-sum test) (Figure 4D).

**Figure 4.**
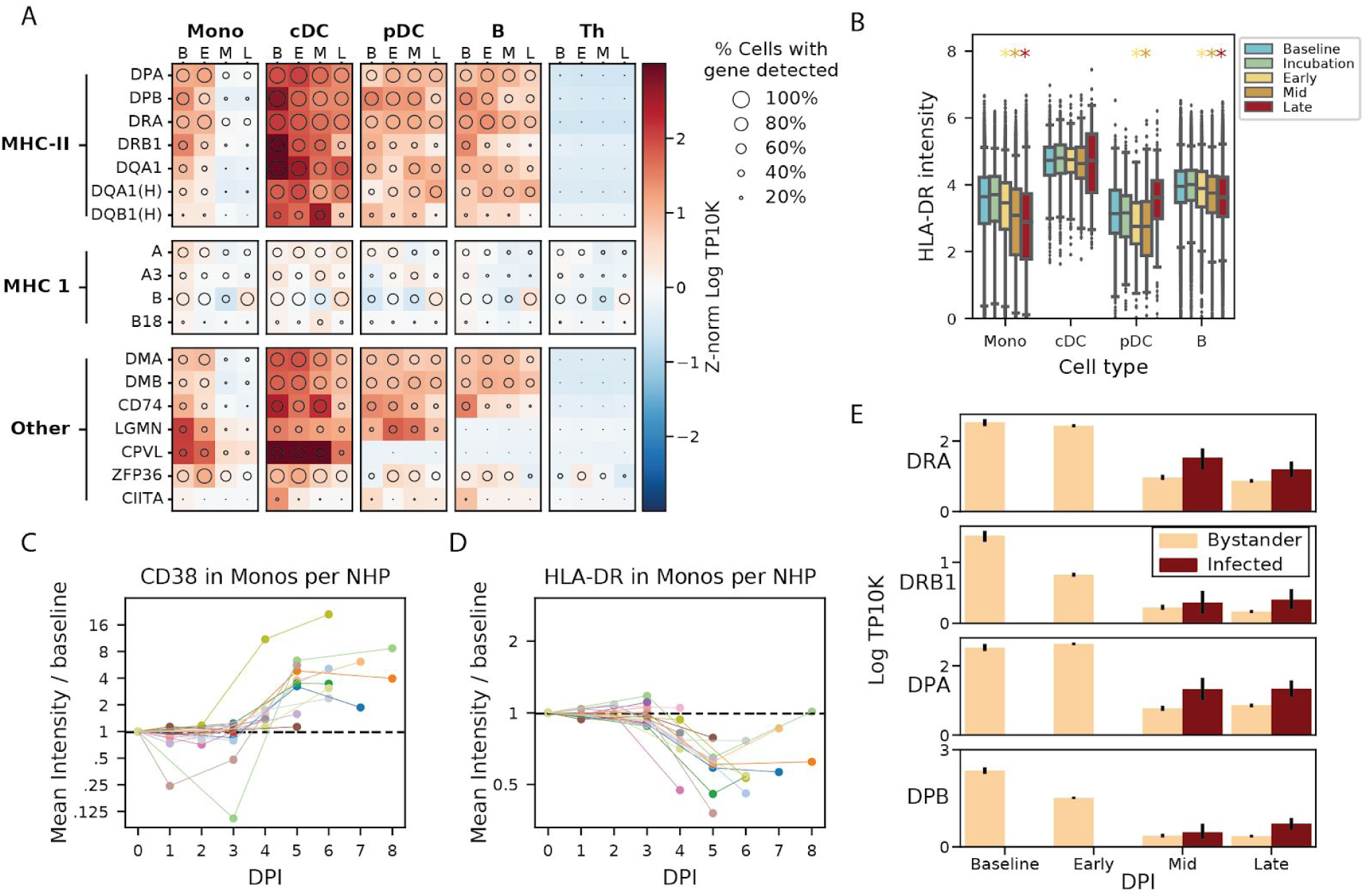
Monocytes dramatically reduce expression of MHC class II proteins independent of infection status. (**A**) Expression profiles of major histocompatibility (MHC) or MHC-associated genes (rows) in key cell types at baseline (B), early (E), middle (M), or late (L) stages of disease (columns). Circle size represents the percentage of cells in that group in which the gene was detected, and color denotes the average expression level, in Z-score normalized, natural log transcripts per 10,000 (TP10K). The “MAMU-” prefix, which designates major histocompatibility genes in rhesus monkeys, was removed from all MHC I and II gene symbols. Gene symbols that initially contained an “HLA-” prefix are indicated with an “(H)”. (**B**) CyTOF intensity (arbitrary units) of HLA-DR protein in antigen-presenting cells at different stages of EVD. Boxes denote the median and interquartile range, and whiskers denote the 2.5th and 97.5th percentiles. Colored stars indicate EVD stages that decreased significantly from baseline (rank-sum test *p* < 0.05) with color corresponding to stage. **(C and D)** Fold change (log_2_ scale) in average CD38 (C) and HLA-DR (D) CyTOF intensity on monocytes at each DPI relative to baseline for each PBMC sample. Colored lines connect serial samples from the same NHP. See also Figure S6C. (**E**) Average gene expression (natural log TP10K) for four MHC class II genes in monocytes at different disease stages, stratified by cell infection status. Error bars denote 95% confidence intervals for the mean based on 200 bootstraps.

Reduced MHC-II expression in monocytes was not a direct consequence of EBOV infection. Only a small (∼5%) percentage of monocytes were infected at mid EVD (Figure 2I), suggesting that the striking decreased levels of MHC-II genes was unlikely to be specific to infected cells. Moreover, we confirmed that the average expression of MHC-II genes was comparable or even higher in infected cells relative to uninfected cells in NHPs with EVD (*i.e.* bystanders) (Figure 4E). Thus we conclude that the MHC-II decrease in monocytes observed in EVD is independent of direct viral infection.

To identify co-regulated genes as well as possible drivers of MHC-II down-regulation, we looked for other genes with expression correlated with MHC-II in monocytes (**Materials and Methods**). Many of the most correlated genes were functionally involved in the antigen presentation pathway, such as *CD74* (Spearman ρ = 0.42, *p* = 1.8×10^-296^), which chaperones MHC-II to the endosome and prevents premature binding of antigen (Schröder, 2016); *LGMN* (Spearman ρ = 0.41, *p* = 8.1×10^-286^), a protease that cleaves proteins to facilitate peptide presentation on MHC-II (Dall and Brandstetter, 2016); and, *B2M* (Spearman ρ = 0.33, 2.6×10^-175^), a component of the MHC class I complex (Figure 4A). In addition, one of the most associated genes was *ZFP36* (Spearman ρ = 0.43, *p* < 1×10^-296^), a protein that directly regulates mRNA stability and turnover of MHC-II and other immune-related RNAs (Pisapia et al., 2019). These findings suggest that MHC-II and other genes involved in antigen presentation may be part of a single transcriptional module, co-regulated by *ZFP36* and/or other genes.

### Characterization of differentially expressed genes between infected and bystander monocytes

Next we characterized genes that were differentially expressed between infected and bystander monocytes, as these could represent host entry factors, restriction factors, or genes that are regulated by infection within a cell. For this and all subsequent differential expression analyses, EBOV transcripts were excluded from the denominator when normalizing cells by library size, to avoid a bias in the estimated expression levels of host genes in infected cells (**Materials and Methods**). We identified 505 genes that were differentially expressed between infected and bystander monocytes (*q* < 0.05) of which 276 changed by more than 30% (Figure 5A, **Table S4**). 181 (18%) of the 1,020 genes that were differentially expressed in monocytes at one or more stages of EVD were also differentially expressed in infected monocytes relative to bystanders.

**Figure 5.**
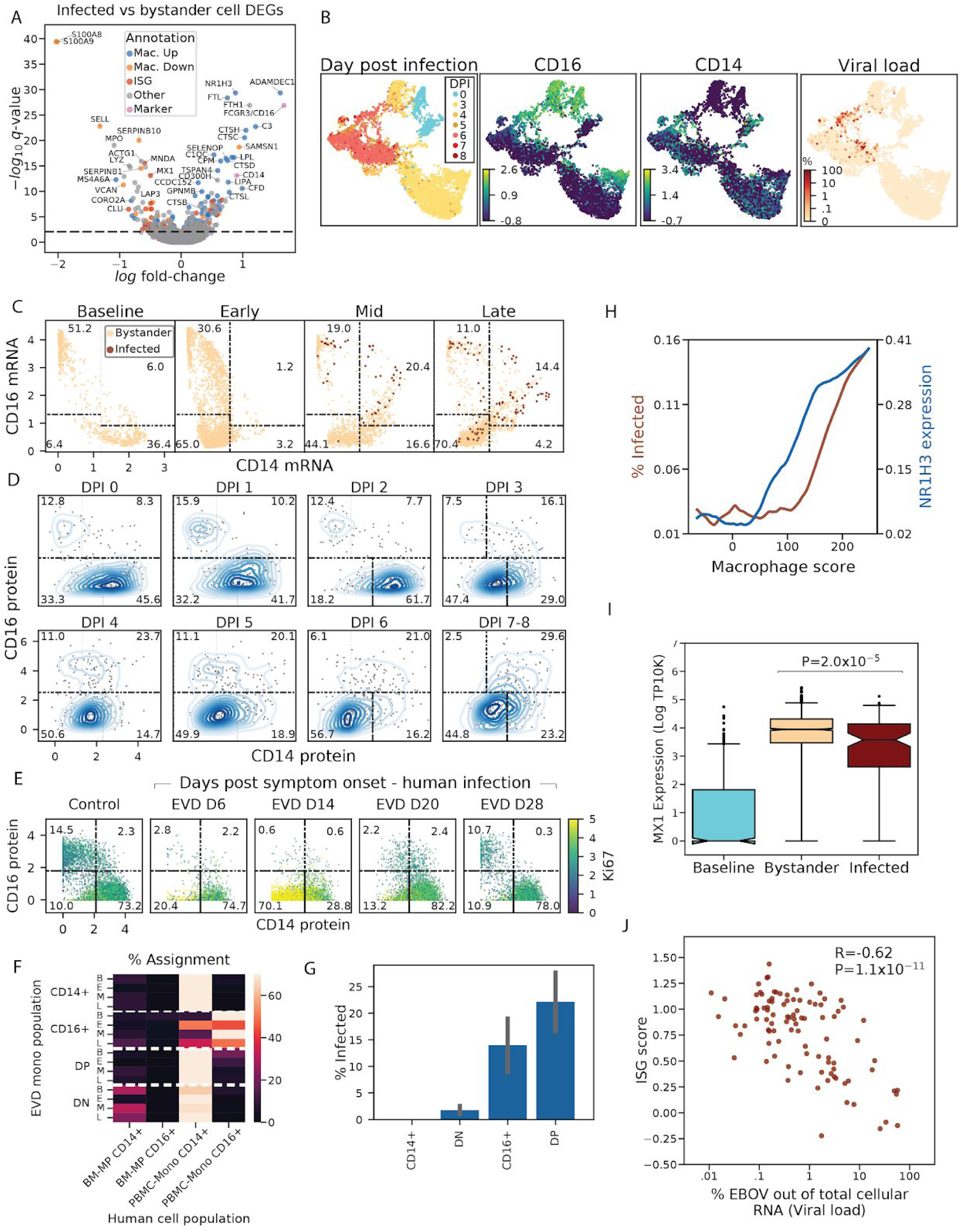
ISG suppression, co-expression of CD14 and CD16, and expression of macrophage genes, are associated with monocyte infectivity. (**A**) Volcano plot of differential expression between infected and bystander monocytes from DPI 5–8. Genes are colored by membership in sets of genes up- (Mac. Up, blue) or down-regulated (Mac. Down, orange) during *in vitro* differentiation of monocytes into macrophages, interferon stimulated genes (ISG, red), or the marker genes *CD14* and *CD16* (Marker, pink). See also **Table S4**. (**B**) UMAP embedding of monocyte gene expression data, colored by (left-to-right): Day post infection, *CD16* expression (log TP10K), *CD14* expression (log TP10K), and percentage of cellular transcripts mapping to EBOV. (**C**) Scatter plot of smoothed expression (log TP10K) of *CD14* and *CD16* for monocytes in baseline, early, mid, and late disease. Cells are colored by infection status. Boxes define the CD14+, CD16+, DN, and DP subsets described in the text, and numbers denote the percentage of cells in each subset at that disease stage. See also Figures S7A and S7B. (**D**) CD14 and CD16 protein (CyTOF intensity) expression on monocytes at each DPI. The distribution is displayed as a bivariate kernel density plot with 200 randomly sampled cells overlaid as a scatter plot. See also Figure S7C. (**E**) CD14 and CD16 protein (CyTOF intensity) expression on monocytes in a case of human EVD, colored by Ki67 protein expression (CyTOF intensity) for multiple days post symptom onset. See also Figure S7D. (**F**) Percent assignment of NHP CD14/CD16 subsets at each EVD stage (B - baseline, E - early, M - mid, L - Late) to human myeloid populations in the reference dataset (BM-MP - bone marrow monocyte progenitors, PBMC-CD16+ - circulating CD16+ monocytes, PBMC-CD14+ - circulating CD14+ monocytes). See also Figures S7E–S7K. (**G**) Percentage of infected monocytes in each CD14/CD16 subset in late infection. Error bars denote 95% confidence intervals for the mean based on 1,000 bootstraps. (**H**) Association between macrophage score (*x*-axis) and percentage of infected cells (left *y*-axis, red), and expression of *NR1H3* (natural log TP10K), a marker of macrophage differentiation (right *y*-axis, blue). Monocytes from late disease are ordered by macrophage score, and we averaged percent of infected cells and *NR1H3* expression within a 400-cell sliding window. See also Figures S8A–S8C. (**I**) *MX1* expression (natural log TP10K) in monocytes at baseline, and uninfected bystanders or infected cells in late infection. Boxes denote the median and interquartile range, and whiskers denote the 2.5th and 97.5th percentiles. See also Figure S8D. (**J**) Scatter plot of ISG score (*y*-axis) versus percentage of cellular transcripts mapping to EBOV (*x*-axis) for infected monocytes in late EVD (DPI 6–8).

We observed that the differentially expressed genes fell into 3 broad categories--genes associated with monocyte subtypes, genes associated with monocyte-to-macrophage differentiation, and interferon stimulated genes (ISGs)--which we explore in the sub-sections below.

#### Emergence of CD14- CD16- immature monocyte precursors suggests emergency myelopoiesis

A notable feature of the differential expression profile was that *CD14* and *FCGR3* (which codes for *CD16)* were over-expressed in infected monocytes relative to bystanders. These two genes define classical (CD14+) and non-classical (CD16+) monocytes respectively (Kapellos et al., 2019), which comprise the dominant monocyte subsets in the blood of healthy individuals.

Classical monocytes are highly phagocytic scavenger cells, while non-classical monocytes are involved in complement and antibody-mediated phagocytosis. To understand the role of monocyte subsets and other genes affecting monocyte heterogeneity across EVD, we visualized the transcriptome profiles of just the monocytes in two dimensions using UMAP. The monocytes separated by DPI and by EBOV infection status, consistent with changing cell states over the disease course (Figure 5B). At baseline, monocytes separated into 2 distinct clusters, marked by high expression of *CD14* (Figure 5B, bottom half of the blue lobe of DPI panel) or *CD16* (Figure 5B, top half of the blue lobe of DPI panel), consistent with the conventional subtyping.

However, monocyte subsets changed dramatically during EVD, with the decline of single-positive CD14+ and CD16+ monocytes, and the corresponding rise of 2 unusual populations: a large population of CD14- CD16- cells (double negatives [DNs]) and a smaller population of CD14+ CD16+ cells (double positives [DPs]). To visualize the dynamics of these populations, we plotted smoothed gene expression of *CD14* and *CD16* for each EVD stage (Figure 5C, **Materials and Methods**). While 87.6% of cells fell into single CD14+ or CD16+ bins at baseline, this dropped to 33.8%, 35.6%, and 15.2% in the early, middle, and late stages of EVD, respectively. We confirmed a corresponding loss of single-positive CD14+ or CD16+ monocytes and a gain of DNs and DPs at the protein level by CyTOF (Figure 5D). In both RNA and protein measurements, these two populations began declining on DPI 3.

At late EVD, the most frequent monocyte population was CD14- CD16- DN cells, which rose to make up 70.4% (by scRNA-Seq) or 56.7% (by CyTOF) of the monocytes. Given that they expressed neither of the canonical (CD14/CD16) monocyte marker genes, we confirmed that their overall gene expression profiles were most correlated with single-positive monocyte populations in late EVD (Pearson correlation R=0.80 CD14+, R=0.58 CD16+) and CD14+ monocytes at baseline (R=0.40), and were less correlated with neutrophils, DCs, and lymphocytes in late EVD (R=0.28, R=0.39, R=0.14, respectively) (Figure S7A). DNs first emerged on DPI 3, coinciding with the 2-fold increase in monocytes we observed on that day (Figure 2D). The DN population is highly proliferative; while 0% of monocytes at baseline expressed moderate levels of Ki67 (smoothed log TP10K > 1), over 37% of DN monocytes expressed Ki67 beyond this threshold by late EVD (Figures S7B **and** S7C). Therefore, this population underlies the increased monocyte replication rates observed in the middle stage of EVD (Figure 2G).

To determine if a corresponding increase in DNs occurs in acute EVD in humans, we re-analyzed published CyTOF data from the 2013–2015 outbreak of 4 acute EVD cases that were treated at the Emory University Hospital Serious Communicable Diseases Unit (McElroy et al., 2020) (**Materials and Methods**). All of the human cases showed a congruent pattern to the NHP data, with loss of conventional CD14+ and CD16+ single-positive monocytes and an emergence of proliferative (Ki67^hi^) DN monocytes (Figures 5E **and** S7D). Then, the DNs disappeared and were replaced by the conventional CD14+ and CD16+ single positive monocytes at later days post symptom onset, as the human cases entered the convalescent period. Thus, the emergence of circulating DNs and the loss of conventional circulating monocyte subsets is a feature of human clinical cases as well as our NHP model of lethal EVD.

The presence of proliferating DN monocytes was surprising because mature monocytes in circulation are believed to be non-dividing (van Furth et al., 1979). However, infectious and neoplastic diseases produce cytokines such as M-CSF that induce the release of proliferating immature myeloid cells from the bone marrow, a process known as emergency myelopoiesis (Chiba et al., 2018; Cuenca et al., 2015; Sayed et al., 2019). We therefore hypothesized that the DN population may reflect immature myeloid cells released from the bone marrow by emergency myelopoiesis.

If DNs represent the product of emergency myelopoiesis, we might expect their gene expression profiles to be more similar to bone marrow resident monocyte precursors than circulating monocytes. To test this, we compared our monocyte populations against a reference scRNASeq dataset of bone marrow monocyte precursors (BM-MPs) from healthy human bone marrow (Hay et al., 2018) and mature monocytes from human PBMCs (Figures S7E–H, **Materials and Methods**). Notably, the BM-MPs showed lower expression of *CD14* and *FCGR3A* (the human *CD16* gene) than mature monocytes, consistent with the diminished expression of these genes in DNs relative to baseline single-positive monocytes in the NHP data (Figure S7I). In addition, the BM-MPs showed higher expression of *MPO*, *AZU1*, *S100A8*, and *S100A9* than mature monocytes (Figures S7I **and** S7J), consistent with the observation that these genes are expressed at higher levels in DNs relative to baseline monocytes (Figure S7K). To formally test whether our monocyte populations were more similar to the mature PBMCs or the BM-MPs, we identified the nearest neighbor of each NHP monocyte in the reference dataset. As expected, DNs from mid and late EVD were significantly more likely to be matched with BM-MPs (32% at mid, 23% late), than single positive CD14+ or CD16+ monocytes, which almost exclusively were assigned to the corresponding circulating monocyte populations in the reference data (<10% assignment to BM-MPs assignment to BM-MPs for all non-DN populations and stages, except for CD16+ at late EVD which was 15%) (Figure 5F).These findings suggest that DNs represent immature monocytes released from the bone marrow in response to the EVD cytokine milieu.

#### Monocytes expressing markers of macrophage differentiation are enriched for EBOV infection

In addition to DNs, we also observed CD14+ CD16+ DP cells, which rose to make up 20.4% of the monocytes by mid EVD (Figure 5C). A similar increase in DP monocytes has been observed in sepsis (Fingerle et al., 1993; Nockher and Scherberich, 1998) and other viral infections (Michlmayr et al., 2018; Zanini et al., 2018a). We found that the DP population harbored a disproportionately high percentage of EBOV-infected cells (Figure 5G), consistent with the fact that *CD14* and *CD16* were both independently higher in EBOV infected monocytes than in bystanders (Figure 5A). At the late infection timepoints, 22.1% of DPs were infected compared to only 1.74% of DNs. Thus, the differential expression of *CD14* and *CD16* in infected cells results from increased infection of the DP cells, rather than increased expression of *CD14* on classical and *CD16* on non-classical monocytes.

We noticed that the differentially expressed genes between infected and bystander monocytes (Figure 5A, **Table S4**) *and* between DP and DN monocytes (Figure S8A, **Table S5**), were enriched for monocyte-to-macrophage differentiation associated genes, including known EBOV entry factors. It has been previously observed that freshly isolated monocytes are largely refractory to EBOV infection in cell culture, but that EBOV entry factors are up-regulated during *in vitro* macrophage differentiation, allowing increased infection (Martinez et al., 2013). *In vivo*, we observed higher levels of macrophage differentiation markers such as *NR1H3*, *ADAMDEC1*, and several cathepsins in infected cells relative to bystanders. Among these genes, the known EBOV entry factors cathepsin L (*CTSL*) and B (*CTSB*), and *GNPTAB* (Carette et al., 2011; Gnirß et al., 2012) were all expressed at significantly higher levels in infected cells than bystanders (*q* = 6.7×10^-9^, 3.8×10^-7^, and 2.1×10^-3^, respectively). By contrast, the cellular receptor *NPC1* was not significantly differentially expressed, suggesting that natural variability in the mRNA abundance of *NPC1* likely does not influence EBOV infectivity within circulating monocytes in rhesus monkeys.

We suspected that up-regulation of the entry factors *CTSL*, *CTSB*, and *GNPTAB* might be occurring as part of a general macrophage differentiation program. We tested this hypothesis using gene sets derived from published bulk RNA-Seq data of primary blood monocytes before and after differentiation into macrophages, *in vitro* (Dong et al., 2013) (**Table S4**). We found that genes that are up-regulated during *in vitro* differentiation were significantly enriched in infected cells (OR = 3.5, *p* = 3.1×10^-11^, Fisher’s exact test) and genes that were down-regulated during differentiation were significantly enriched in bystanders (OR = 3.7, *p* = 4.2×10^-8^, Fisher’s exact test; combined chi-squared goodness of fit test *p* = 2.2×10^-30^). Using gene set annotations from two other RNA-Seq studies of macrophage *in vitro* differentiation resulted in similar findings (chi-squared goodness of fit test *p* = 2.6×10^-9 (^Saeed et al., 2014^)^, Fisher’s exact test *p* = 5.7×10^-12 (^Italiani et al., 2014^)^). Genes associated with differentiation into M2-polarized macrophages were more enriched among EBOV-infected cells than those of M1-polarized macrophages (OR = 7.8, *p* = 1.3×10^-10^ compared to OR = 3.3, *p* = 1.6×10^-3 (^Italiani et al., 2014^)^, **Table S4**).

Next, we quantified the proportion of infected cells as a function of macrophage-differentiation, and found that infectivity increased along with expression of the macrophage differentiation program. To determine the relative activity of the macrophage program in each cell, we computed a “macrophage differentiation score” consisting of a weighted sum of the 618 genes that were significantly positively or negatively correlated with *in vitro* differentiation in the (Dong et al., 2013) dataset (**Materials and Methods**). Ranking cells from lowest to highest macrophage score, we observed that the percentage of infected cells rose more than four-fold from 3.0% to 15.0% (Figure 5H). Thus, our data strongly suggests that of all circulating cells, EBOV predominantly infects monocytes with the highest expression of the macrophage differentiation program.

Given that macrophage differentiation genes were over-expressed in DPs relative to DNs (Figure S8A), we sought to understand the relationship between the CD14 and CD16 defined monocyte subsets and expression of the macrophage differentiation program. Comparing the overall macrophage score of the different CD14/CD16-marked subpopulations confirmed that DPs generally had the highest macrophage scores while DNs and single CD14+ cells had the lowest (Figure S8B). Thus, some of the enrichment of infected cells among the DP subset could potentially be attributed to their more ‘macrophage-like’ gene expression.

However, there was substantial heterogeneity in macrophage scores within DPs and the other CD14/CD16-marked subsets, and we found that macrophage score and CD14/CD16 subset were independently predictive of infectivity. To demonstrate this, we stratified cells in each subset by macrophage score (above or below the median value across all subsets combined). This showed that less macrophage-like DPs were still more likely to be infected than more macrophage-like DNs, even though all cells in the former category had a lower macrophage score than all cells in the latter (*p* =1.9×10^-25^, Fisher’s exact Test, Figure S8C). However, within the DPs, more macrophage-like cells were more likely to be infected than less macrophage-like cells (*p* = 0.0003, Fisher’s exact Test, Figure S8C). This suggests that infection could not be explained by either CD14/CD16 subset or macrophage score alone. As a further confirmation, we fit a logistic regression predicting the infection status of each cell using macrophage score, smoothed *CD14* and *CD16* expression values, and a *CD14*x*CD16* interaction term (**Materials and Methods**). As expected, the *CD14*x*CD16* interaction term (which is highest in DPs) and macrophage score were positively associated with infection status, and the *CD14* and *CD16* terms were negatively associated (*p* < 0.01 for all coefficients). These findings demonstrate that monocyte CD14/CD16 subset and differentiation status independently impact the probability of a cell being infected with EBOV, *in vivo*.

#### Interferon stimulated genes are down-regulated in infected monocytes relative to bystanders

Finally, we noticed that several key ISGs such as *MX1* were expressed at lower levels in infected cells than in bystanders (MAST *q* = 7.7×10^-14^, rank-sum test *p* = 2.0×10^-5^, Figures 5A and 5I). To determine if infection had a suppressive effect on overall ISG expression, we compared the magnitude of the interferon response (defined previously as the ISG score, **Materials and Methods**) between infected and bystander cells at late EVD. While both bystander and infected monocytes at late EVD had higher ISG scores than monocytes at baseline, ISG scores were lower in infected cells than bystanders (not statistically significant by rank-sum test, Figure S8D). More strikingly, there was a significant negative correlation between ISG score and the percentage of cellular transcripts derived from EBOV (*i.e.*, the intracellular viral load) (Spearman ρ = −0.62, *p* = 1.1×10^-11^, Figure 5J). This suggests that ISGs are down-regulated during viral replication within infected cells (see Figure 7 and **Discussion**).

### Single-cell transcriptomics of *ex vivo* infected PBMCs reveals temporal dynamics in viral gene expression

In order to more thoroughly probe viral and host gene expression changes during the viral life cycle, we sought to obtain transcriptomes from a greater number of infected cells. Thus, we isolated PBMCs from 2 healthy rhesus monkeys (NHP1 and NHP2) and inoculated them *ex vivo* with either live EBOV, EBOV rendered replication-incompetent by gamma irradiation (Feldmann et al., 2019), or media only as a control (Figure 6A). We selected a multiplicity of infection (MOI) of 0.1 plaque forming units (pfu, titrated on Vero E6 cells)/cell to ensure that a large proportion of cells would be infected. We performed scRNA-Seq using Seq-Well at 4 hours or 24 hours post-infection (HPI), corresponding to very early (start of viral transcription) and middle-to-late stages (viral genome replication, virion assembly) of the viral life cycle. Inoculation with gamma-irradiated EBOV allowed us to characterize the host response in the absence of effective viral transcription and translation.

**Figure 6.**
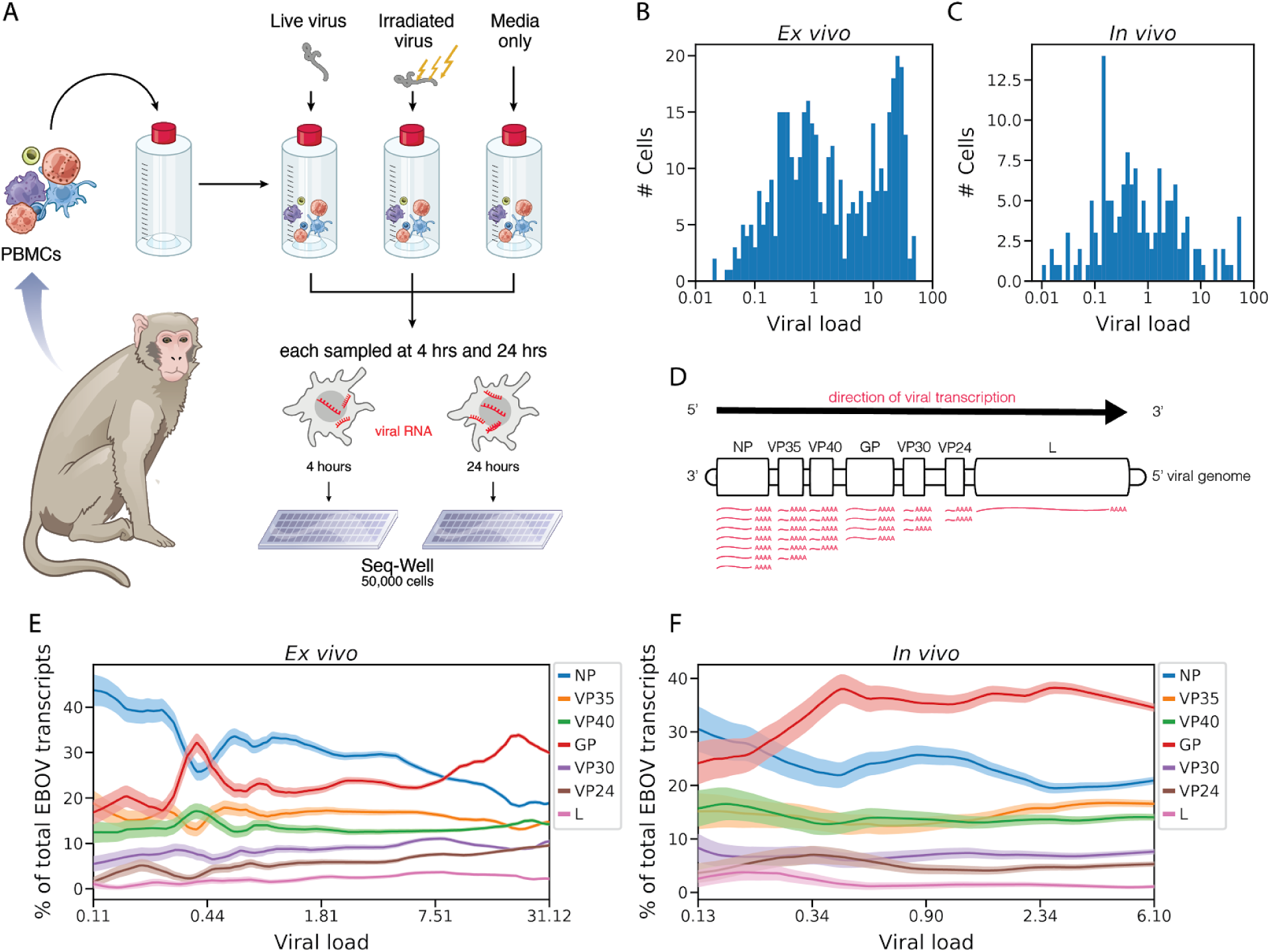
Viral transcriptional dynamics of infected monocytes *in vivo* and *ex vivo*. (**A**) Schematic of EBOV challenge of PBMCs *ex vivo*. See also Figure S9. (**B and C**) Histogram of the percentage of cellular transcripts derived from EBOV (intracellular viral load) in monocytes from PBMCs inoculated with live virus *ex vivo* (B) or from PBMCs of NHPs infected *in vivo* (C). See also Figures S10A–S10D. (**D**) Schematic of EBOV transcription. The viral RNA-directed RNA-polymerase transcribes each gene sequentially from the 19 kb genome, but occasionally releases the genomic RNA template thereby ending transcription. As a result, *NP* is transcribed most frequently, and *L* least frequently. (**E and F**) Relative proportion of each EBOV gene versus viral load (log_10_ scale), *ex vivo* (E) or *in vivo* (F). All infected monocytes were ordered by viral load and the percentage of each viral gene was averaged over a 50-cell sliding window. Bands denote the mean ± 1 SD. See also Figures S10E and S10F.

**Figure 7.**
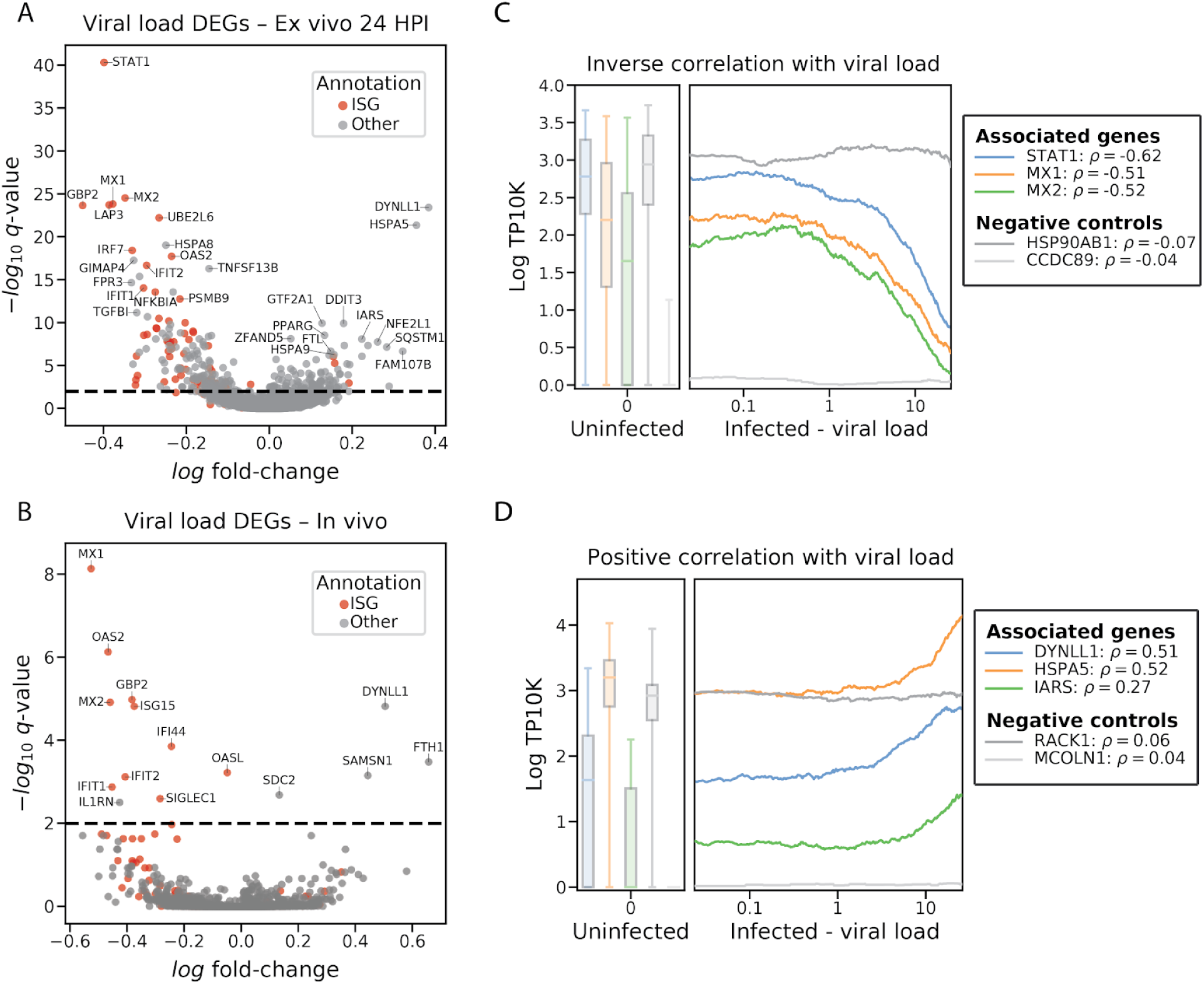
EBOV infection down-regulates host antiviral genes and up-regulates putative pro-viral genes. (**A and B**) Volcano plot of association between host genes expression and viral load, within infected monocytes from PBMCs 24 HPI treated with live virus *ex vivo* (A) or from PBMCs of NHPs infected *in vivo* on DPI 5-8 (B). See also **Table S6**. (**C and D**) Association between host gene expression and viral load for selected positively (C) and negatively (D) associated genes in monocytes from *ex vivo* infections. Infected cells are ordered by viral load (log_10_ scale), and we depict gene expression (natural log TP10K) averaged over a 100-cell sliding window. Genes that are significantly associated with viral load are shown in color while unassociated negative control genes are shown in gray. Spearman correlation coefficients (ρ) for the association between viral load and gene expression are listed in the legend. Box plots represent gene expression in uninfected cells (boxes denote the median and interquartile range, and whiskers denote the 2.5th and 97.5th percentiles) See also Figures S10G and S10H.

We obtained single-cell transcriptomes from 50,646 PBMCs inoculated *ex vivo*, and observed similar cell-type representation, clustering by treatment condition, and distribution of EBOV-infected cells as with the *in vivo* collections (Figures S9A–S9C), with a few notable exceptions. First, we observed that cells from NHP1 and NHP2 separated in UMAP embeddings (Figure S9D) and that this separation was associated with higher expression of ISGs such as *MX1* in cells derived from NHP1 compared to NHP2 (Figures S9E **and** S9F). The ISG signal was most predominant in cells from NHP1 at 24 HPI treated with either irradiated or live virus (Figure S9G). We therefore analyzed cells from each animal both separately and jointly to avoid potential artifacts. A second difference is that while we did not observe infected DCs *in vivo*, we found that 16% of DCs (16.0% in NHP1, 15.7% in NHP2) inoculated with live virus *ex vivo* were infected by 24 HPI (Figure S9H). This difference could be due to increased density of cells in culture, the higher effective MOI we used in the *ex vivo* experiment, or even changes to the expression states of DCs associated with culture conditions.

Consistent with the *in vivo* data, monocytes were the predominant infected cell type, with over 65% infected by 24 HPI after inoculation with live virus (76% in NHP1, 61% in NHP2) (Figures S9H **and** S9I). 11.8% of the monocytes treated with irradiated virus also contained a statistically significant number of viral reads by 24 HPI, despite the fact that gamma irradiation induces damage to the viral genome that eliminates productive viral replication (Feldmann et al., 2019). As expected, cells treated with irradiated virus had a significantly lower fraction of EBOV reads per cell than those treated with live virus (Figure S10A). Moreover, viral RNAs from the cells treated with irradiated virus were substantially less likely to be coding-sense mRNA transcripts than anti-sense viral genomic RNA. For live-virus treated cells, 78% and 92% of detected RNAs were mRNA at 4 and 24 HPI, compared to only 37% and 44% in cells treated with irradiated virus (Figure S10B). This suggests that our method can detect fragments of viral genomic RNA from irradiated virus that have entered cells, but as expected, these do not reflect productive infections and do not generate significant quantities of poly-adenylated mRNAs.

Exploiting the increased resolution of the *ex vivo* dataset, we characterized the heterogeneity in viral transcript abundance per cell (*i.e.*, the intracellular viral load). We observed that the intracellular viral load varies over several orders of magnitude in infected cells, both *in vivo* and *ex vivo* (Figures 6B **and** 6C). While most cells harbored viral loads below 0.1%, a substantial minority had loads of >10%, with maximum detected loads of 57.5% and 52.3% for cells *in vivo* and *ex vivo* respectively. The observed heterogeneity in viral load was not due to different numbers of transcripts detected per cell, because cells with low and high viral load had a similar range of total transcripts detected (Figures S10C **and** S10D).

We next analyzed the dynamics of EBOV gene expression to determine if it matched the predicted pattern based on established models of EBOV transcription. Transcription of EBOV’s 7 genes by the viral RNA-directed RNA polymerase L follows the canonical stop-start mechanism described for filoviruses and other non-segmented negative-strand RNA viruses (Brauburger et al., 2014, 2016). L initiates transcription *de novo* (Deflubé et al., 2019) at the 3’ end of the genome, and processes from 5’ to 3’; at each gene transcription termination signal, L pauses and either falls off the genomic RNA template or reinitiates transcription of a new mRNA for the subsequent gene (Figure 6D, (Mühlberger, 2007)). As a consequence, *NP* is the first gene to be transcribed and is transcribed at the highest level, proceeding down the genome to the polymerase gene *L* being transcribed last and at the lowest level.

When we quantified the relative expression levels of EBOV genes as a function of viral load, we observed an unexpected accumulation of *GP* mRNA (Figures 6E **and** 6F) that was consistent between NHP1 and NHP2 (Figures S10E **and** S10F). At low viral loads, both *in vivo* and *ex vivo*, the gene expression distribution roughly matched the expected pattern, with most of the transcripts derived from the 3’ end of the genome, in particular *NP*, and the fewest transcripts derived from the 5’ genes *VP30*, *VP24*, and *L*. In agreement with this pattern, cells inoculated with irradiated virus, which has impaired transcription due to RNA cleavage or crosslinking (Feng et al., 2011; Ginoza, 1967; Ward, 1980), were highly enriched in *NP* mRNA (Figure S10B), suggestive of RNA fragment transcription. However, as viral load increased in cells infected with live EBOV, *GP* was the most highly expressed viral transcript. This finding is unexpected based on the start-stop mechanism where *NP* should be transcribed at a strictly higher rate than *GP*. This observation suggests a life-cycle dependent regulation of viral gene expression that has not previously been observed for EBOV (see **Discussion**).

### EBOV infection down-regulates host antiviral genes and up-regulates putative pro-viral genes

Next, we exploited natural variability in viral load across infected cells to identify host gene expression changes correlated with viral replication, which may therefore represent pathways directly regulated by infection. Instead of testing for differential expression between infected and bystander cells as we did previously to define tropism-associated genes, we looked for continuous association between viral abundance and host transcript levels in infected monocytes (**Materials and Methods**). This identified 264 genes that were negatively correlated and 211 genes that were positively correlated with viral load *ex vivo* with *q* < 0.05, of which 34 changed by more than 30% per 10-fold increase in viral load (Figure 7A, **Table S6**).

Consistent with our previous observation that ISG score decreased with viral load within a cell, ISGs decreased dramatically with EBOV levels, both *in vivo* and *ex vivo* (e.g., *MX1*, *q* = 1.5×10^-24^ *ex vivo*, *q* = 7.4×10^-9^ *in vivo*, Figures 7A and 7B, **Table S6**). *Ex vivo*, the most negatively associated gene was *STAT1*, the master transcriptional regulator of the IFN response (*q* = 4.9×10^-41^ *ex vivo*, *p =* 0.0072 *in vivo* but *not significant by FDR*). Previous experiments have shown that the EBOV protein VP24 inhibits STAT1 activity by blocking its translocation to the nucleus (Leung et al., 2006) through direct binding (Zhang et al., 2012). However, this is the first observation that *STAT1* mRNA levels decrease with viral replication within infected cells *in vivo*.

We visualized the expression of *STAT1* and several other negatively regulated ISGs over the course of infection within a cell (Figure 7C). The expression of these genes remained relatively constant as EBOV levels rose to 1% of cellular transcripts; however as EBOV levels continued to rise beyond 1%, the expression of *STAT1* and many of its target genes declined precipitously. This suggests that there is a delay before EBOV can transcriptionally down-regulate host antiviral genes since it must transcribe and translate VP24 and other immunomodulatory proteins before they can act. The trajectories of these host antiviral genes were consistent between donor animals *ex vivo* (Figure S10G) despite the fact that more cells from NHP1 mounted an IFN response than NHP2.

Only a handful of host genes increased in expression level alongside viral load, but their trajectories were consistent between the two animals (Figure S10H). The most dramatically up-regulated gene was *DYNLL1* (*q* = 2.5×10^-27^ *ex vivo*, *q* = 1.5×10^-5^ *in vivo*, **Table S6**), which increased significantly both *in vivo* and *ex vivo* (Figures 7A and 7B). DYNLL1 is a multi-functional protein involved in intracellular transport (Barbar, 2008). Intriguingly, DYNLL1 was previously shown to increase EBOV replication in a minigenome reporter assay (Luthra et al., 2015). Our data show that *DYNLL1* mRNA is up-regulated *in vivo* and *ex vivo*, starting when EBOV RNA constitutes between 0.1–1% of the transcripts in a cell, before ISGs begin decreasing (Figure 7D).

Several other genes that we identified as up-regulated alongside viral replication have known or speculative pro-viral functions in protein folding and synthesis. For example, *HSPA5* (*q* = 4.5×10^-22^ *ex vivo*, *p* = 0.04 *in vivo* but *not significant* by FDR) encodes a chaperone protein that was previously shown to be an essential host factor for EBOV (Reid et al., 2014). However, this is the first observation that *HSPA5* mRNA is up-regulated in EBOV infection. Other hits such as *DDIT3* and *NFE2L1* (*q* = 1.3×10^-10^ and *q* = 1.8×10^-8^ respectively*, ex vivo*) are sensors of ER and oxidative stress (Kim et al., 2016) and have been implicated in cell lines infected with Marburg virus (Hölzer et al., 2016) and in monocytes in EBOV-infected NHPs (Menicucci et al., 2017). We observe a corresponding enrichment of gene sets associated with ER stress response among genes up-regulated alongside intracellular viral load such as BUYTAERT_PHOTODYNAMIC_THERAPY_STRESS_UP, PODAR_RESPONSE_TO_ADAPHOSTIN_UP, and HALLMARK_UNFOLDED_PROTEIN_RESPONSE (*q* = 4.7×10^-14^, *q* = 5.3 x 10^-5^, *q* = 0.004 respectively, Fisher’s exact test, **Table S6**). This suggests that oxidative and/or ER stress may play an important role in the pathophysiology of infected cells, particularly at a late stage of viral replication.

Increased translation can also deplete tRNAs and free amino acids (Albers and Czech, 2016). We observed statistically significant up-regulation of *IARS* - isoleucine tRNA synthetase (*q* = 8.4×10^-9^ *ex vivo*) - which may reflect the cellular response to increased translational demand due to overwhelming production of viral proteins. In addition to *IARS*, several gene sets associated with depletion of amino acids were significantly up-regulated such as KRIGE_RESPONSE_TO_TOSEDOSTAT_24HR_UP (Tosedostat is an aminopeptidase inhibitor), KRIGE_AMINO_ACID_DEPRIVATION and PENG_LEUCINE_DEPRIVATION_UP (*q* = 2.3×10^-6^, *q* = 0.0030, *q* = 0.0040, respectively, Fisher’s exact test, **Table S6**), suggesting that viral replication exhausts cellular amino acid stores. This hypothesis is consistent with prior observations of depleted amino acids in the plasma of fatal human EVD cases (Eisfeld et al., 2017).

## Discussion

Despite recurrent outbreaks, the molecular basis of EVD pathogenesis in humans remains understudied due to the biosafety and logistical challenges associated with performing clinical research during outbreaks in resource-poor settings, and experimentally investigating EBOV in maximum containment facilities. By adapting CyTOF and scRNA-Seq approaches for use in BSL-4 containment, we comprehensively surveyed the molecular correlates of disease progression and viral replication in circulating immune cells in a nonhuman primate model of EVD. This study, which is the first high-parameter, single-cell investigation of a RG-4 agent, shed new light on changes in cell-type abundance throughout lethal EVD, defined the preferred targets of EBOV amongst circulating cells, and identified genes regulated by the cytokine milieu or by direct EBOV infection.

We characterized transcriptional- and protein-level changes in monocytes during EVD in NHPs, some of which reflect disruption of their physiological antiviral function. Monocytes had over twice as many differentially expressed genes as other cell types, including genes involved in IFN response, cytokine production, myeloid differentiation, and antigen presentation (Figures 3 **and** 4). Monocytes became activated by IFN during EVD, which normally triggers an increase in expression of multiple MHC-II genes for antigen presentation (Steimle et al., 1994). Surprisingly, almost all of the MHC-II genes and other genes in the antigen presentation pathway were strikingly down-regulated in monocytes, with modest changes in B cells, and no significant changes in cDCs. Moreover, MHC-II genes decreased in both infected and uninfected monocytes, suggesting that the cytokine milieu led to decreased MHC-II expression (Figure 4). These findings suggest a failure of monocytes to present antigens, which might explain why a failed or delayed adaptive immune response is a hallmark of fatal EVD in humans (Baize et al., 1999; Lüdtke et al., 2016).

As EVD progressed, conventional CD14+ and CD16+ monocyte subsets disappeared and were replaced by two unusual populations: CD14+ CD16+ (DP) monocytes, which are known to increase in other infections (Fingerle et al., 1993; Michlmayr et al., 2018; Nockher and Scherberich, 1998; Zanini et al., 2018a), and an unexpected CD14- CD16- (DN) population, that, to our knowledge, has not been previously described in viral infections (Figures 5C–5F). The DN monocytes were highly proliferative and their transcriptomes were more similar to bone marrow resident monocyte precursors than circulating monocytes. This suggests that they may be the product of emergency myelopoiesis, a process whereby cytokines stimulate the bone marrow to release immature myeloid lineage cells that are sometimes referred to as immunosuppressive myeloid cells (Chiba et al., 2018; Hérault et al., 2017; Sayed et al., 2019). DNs had high expression of neutrophil granule genes such as *MPO* and *AZU1*, suggesting that they may represent immature cells prior to the branching of neutrophil and monocyte lineages (*i.e.*, common myeloid progenitors). We identified the emergence of an analogous DN population in human acute EVD cases as well, which later disappeared as the patients recovered, confirming that this population becomes the dominant circulating monocyte population in human disease as well. This finding highlights the power of high-parameter methods such as scRNA-Seq and CyTOF over previous approaches like FACS; despite little-to-no detection of CD14 or CD16, the conventional markers for phenotyping monocytes, there were enough other RNA and protein markers to reliably detect these DN cells as monocyte-lineage cells.

Our data refines the picture of EBOV’s tropism in NHPs, demonstrating that the predominant EBOV infected population in circulation are the DP monocytes which expand during the infection, and monocytes expressing markers of macrophage differentiation (Figures 5C, 5G, **and** 5H). Existing literature has already demonstrated that myeloid cells are major targets of EBOV (Geisbert et al., 2003b, 2003c; Greenberg et al., 2020), including DCs *ex vivo* and in lymph nodes *in vivo* (Geisbert et al., 2003c). While we observe infected monocytes and DCs *ex vivo*, only monocytes were infected more often than expected due to chance among circulating immune cells *in vivo*. This might reflect relevant biological phenomena required for DC infection, such as cell density, cell-to-cell contact, or MOI. Among monocytes, EBOV-infected cells were significantly enriched in CD14+ CD16+ DP monocytes, which were the most macrophage-like and expressed increased differentiation genes, including known EBOV entry factors like cathepsin B (Chandran et al., 2005; Martinez et al., 2013; Schornberg et al., 2006). It has been shown that cultured monocytes are only susceptible to EBOV infection upon differentiation (Martinez et al., 2013), and our data further shows that infectivity *in vivo* strongly correlates with physiological variability of monocyte differentiation state in the context of an active immune system. Furthermore, our data support the hypothesis that the relative abundance of DP monocytes, the preferred circulating cell targets of EBOV, increases over the course of infection, perhaps driven by the cytokine milieu of EVD.

Among infected monocytes, we observed substantial heterogeneity in intracellular viral load, which we exploited as a proxy for staging EBOV’s progression through its life cycle. We confidently identified infected cells with as low as 0.01% EBOV RNA out of total RNA (intracellular viral load), while some cells had up to 57.5% EBOV RNA (Figures 6B **and** 6C). Similar heterogeneity has been observed for the segmented-ssRNA virus influenza (Russell et al., 2018), the +ssRNA virus dengue (Zanini et al., 2018a, 2018b), and the dsDNA virus HCMV (Hein and Weissman, 2019), but this is the first demonstration for a non-segmented negative-sense (NNS) virus.

By analyzing patterns of gene expression among cells at different stages of the viral infection, we identified an unexpected over-representation of *GP* mRNA accumulating late in the viral life cycle. NNS viruses, including EBOV, have strict transcriptional gene regulation based on the viral genome organization (Figure 6D) (Brauburger et al., 2014, 2016). In our data, cells with low viral load (reflecting early stages of the viral life cycle) had relative EBOV transcript abundances that mirrored the genome organization as expected -- *i.e.*, they had the highest expression of *NP* and the lowest expression of *L*. But cells with high viral load (later during the viral life cycle) had higher abundance of *GP* mRNA than *NP* mRNA (Figures 6E and 6F). Given the NNS virus dogma that *NP* is transcribed more frequently than *GP* mRNA, our data suggest that alternate transcription or post-transcriptional regulatory mechanisms, such as increased stability of *GP* mRNA, may account for accumulation of *GP* mRNA late during the viral life cycle. Many viruses increase structural protein production late in the viral life cycle (Honess and Roizman, 1974; Irigoyen et al., 2016; King et al., 2018; Shin et al., 2015); increased EBOV *GP* late in the viral life cycle likely increases the formation of infectious virions.

Many host ISGs negatively correlated with intracellular viral load, suggesting that viral infection down-regulates ISG expression *in vivo* and *ex vivo* (Figures 7A**–C**). These findings strongly suggest that EBOV is specifically down-regulating ISGs rather than preferentially infecting cells with low ISG levels. First, there are multiple well-established mechanisms by which EBOV down-regulates transcription of ISGs, such as by preventing the master antiviral transcription factor, STAT1, from translocating to the nucleus (Harcourt et al., 1999; Kash et al., 2006; Leung et al., 2006; Reid et al., 2008). Second, the percentage of EBOV-infected cells increases gradually from DPI 4 onward (Figure 2I) despite the fact that monocytes are mounting a strong ISG response by then, which suggests that EBOV is able to overcome the inhibitory activities of ISGs and continue replicating. Third, when EBOV infects a cell, viral load must start at a low level before increasing with replication and transcription. If the causal direction was reversed -- *i.e.*, low ISG levels supported EBOV progression and high ISG levels inhibited EBOV progression -- we would expect to see some cells with low ISG and low viral load, but we do not observe any cells with low (0.01–0.1%) viral load and low ISG levels (Figure 5J). Thus our data suggests that EBOV is able to infect monocytes that are mounting a full IFN response, overcome the inhibitory activities of ISGs, and transcribe viral mRNA to high levels during disease, *in vivo*.

On the other hand, several putative pro-viral host genes were positively correlated with intracellular viral load, suggesting that they are directly responding to, or are regulated by, the presence of virus in infected cells (Figures 7A, **7B, and 7D**). *DYNLL1* was the top associated gene with viral load, both *in vivo* and *ex vivo*. Previous work identified an interaction between DYNLL1 and EBOV VP35 (Kubota et al., 2009) that increased EBOV RNA synthesis in a minigenome assay (Luthra et al., 2015). DYNLL1 associates with proteins of many viruses (Merino-Gracia et al., 2011), and increases replication of rabies virus (Tan et al., 2007) an NNS virus similar to EBOV. Here, we showed that *DYNLL1* expression is up-regulated within infected cells *in vivo*, suggesting that EBOV manipulates cellular pathways to encourage a pro-viral cellular environment. Nuclear DYNLL1 typically represses its own transcription factor ATMIN (Jurado et al., 2012); we hypothesize that EBOV VP35 may sequester DYNLL1 protein in the cytoplasm, relieving repression of ATMIN, thus up-regulating *DYNLL1* mRNA.

Additional intriguing genes that are positively correlated with intracellular viral load include *HSPA5*, *NFE2L1*, *DDIT3*, *GTF2A1*, and *IARS*, though their effect sizes are larger *ex vivo* than *in vivo*. Many of these genes are involved in the cellular stress response, which can be triggered when infection overwhelms host translation. As a chaperone protein, *HSPA5* in particular is essential for EBOV replication (Reid et al., 2014); *NFE2L1* and *DDIT3* sense ER and oxidative stress (Kim et al., 2016) and are upregulated upon filovirus infection (Hölzer et al., 2016; Menicucci et al., 2017), and our data suggest that these genes are up-regulated throughout the viral life cycle. Viral transcription and translation can also deplete tRNAs and free amino acids; as a result, viruses have evolved mechanisms to maintain translational capacity (Albers and Czech, 2016). The tRNA synthetase *IARS* could be up-regulated in response to this depleted host environment, also reflected in fatal EVD cases (Eisfeld et al., 2017). By computationally staging cells by their phase in the infection cycle, we nominated several putative pro-viral genes for further study, highlighting the utility of single-cell profiling to study host-virus interactions.

The accumulation of additional host-pathogen single-cell datasets promises to greatly enhance our understanding of infection by allowing us to determine which features of pathogenesis are shared between, or specific to, individual pathogens. For example, our scRNA-Seq and CyTOF data identified several molecular commonalities between EVD and immunosuppressive septic shock (Bray and Mahanty, 2003), which is also characterized by loss of MHC-II expression in monocytes (Monneret and Venet, 2014; Reyes et al., 2020), increased DP monocytes (Fingerle et al., 1993), and emergency myelopoiesis (Bomans et al., 2018; Cuenca et al., 2015; Reyes et al., 2020). Soluble mediators, including cytokines and glucocorticoids, could be key drivers of both EVD and sepsis pathophysiology. TNFα signaling has been extensively implicated as a driver of systemic loss of vascular resistance and shock during EVD. Glucocorticoids have been less well studied in EVD, but decrease MHC-II *ex vivo* (Hawrylowicz et al., 1994) and *in vivo* during sepsis (Tulzo et al., 2004) and reduce *CD14* expression (Nockher and Scherberich, 1997) while increasing the abundance of DP monocytes (Liu et al., 2015). Indeed, the connection between EVD and sepsis may be direct: studies have found bacterial invasion during EVD in NHPs (Reisler et al., 2018) and in humans (Carroll et al., 2017), with immune signatures that resemble sepsis (Eisfeld et al., 2017).

In summary, this work expands our understanding of EVD, and provides a general paradigm for exploring molecular features of host-pathogen interactions, such as tropism and dysregulation of cell circuitry in infected cells, that can be applied to other emerging pathogens.

## Materials and Methods

### Resource Availability

#### Lead Contact

Further information and requests for resources and reagents should be directed to Aaron Lin.

#### Materials Availability

This study did not generate new unique reagents.

### Data and Code Availability

The analysis scripts used in this study are available at https://github.com/dylkot/SC-Ebola.

Datasets generated during this study will be available on GEO and raw sequence data will be available on SRA.

CyTOF .fcs files will be made available via FlowRepository.

### Experimental Model and Subject Details

This study included a subset (21 of 27) outbred rhesus monkeys (*Macaca mulatta*) of Chinese origin described recently (Bennett et al. in submission) (Greenberg et al., 2020). These 27 nonhuman primates (NHPs) were randomized into cohorts (Figure S1B, (Bennett et al. in submission)), balancing age, weight, and sex across 7 groups. All work was approved and performed in accordance with the Guide for the Care and Use of Laboratory Animals of the National Institute of Health, the Office of Animal Welfare, and the US Department of Agriculture (Bennett et al. in submission).

### Method Details

#### Serial sampling study

This study utilized the Ebola virus/H. sapiens-tc/COD/1995/Kikwit-9510621 (EBOV/Kikwit; GenBank accession MG572235.1; *Filoviridae: Zaire ebolavirus*) isolate for the *in vivo* and *ex vivo* challenges, obtained from the Biodefense and Emerging Infections Research Resources Repository (BEI Resources, Manassas, VA, USA). It is the standard challenge stock defined by the filovirus animal non-clinical group (FANG) for testing product efficacy for FDA approval and is well characterized.

For all 21 outbred rhesus monkeys, two baseline blood samples were collected between 0–14 and 14–30 days prior to infection (Figure S1B). 18 NHPs were exposed to the EBOV/Kikwit isolate diluted to a target concentration of 1,000 plaque forming units (PFU) in a volume of 1 mL/dose. All NHPs were inoculated within a 5 month period. This same cohort has already been described recently (Bennett et al. in submission) (Greenberg et al., 2020).

#### Clinical observations and scoring

Beginning on day post-infection (DPI) 0, NHPs were observed 1–3 times daily and given a clinical score based on five criteria: overall clinical appearance and signs of hemorrhage; respiratory rate, mucous membrane color, and dyspnea; recumbency; non-responsiveness; and core temperature (Bennett et al. in submission). Each criterium was assigned a score between 1 and 10, and all scores were added together. Once an NHP reached a combined score of >10, the animal was humanely euthanized.

#### Whole blood collection

Blood was drawn from anesthetized animals into BD vacutainer plastic serum separator tubes (SST) for serum viral load quantification, or in BD vacutainer plastic blood collection tubes with K_3_EDTA for hematology and peripheral blood mononuclear cell (PBMC) purification (Becton Dickinson, Franklin Lakes, NJ, USA) (Bennett et al. in submission). SST tubes were centrifuged at room temperature for 10 minutes (min) at 1800 x *g* to isolate serum. K_3_EDTA tubes were mixed by gentle inversion prior to hematology and PBMC purification.

#### Hematology and complete blood counts (CBC)

250 μL of each whole blood sample was analyzed on a Sysmex 2000i XT (Sysmex Corporation, Kobe, Hyogo Prefecture, Japan) (Bennett et al. in submission). Parameters analyzed by this instrument were: counts of basophils, eosinophils, lymphocytes, monocytes, neutrophils, white blood cell count; percentages of each cell type; and mean platelet volume.

To estimate the abundance of lymphocyte cell types, we multiplied the CBC lymphocyte count by the proportion of lymphocytes of each cell type (CD8 T cells, CD4 T cells, NK cells, and B cells) which was calculated from the unsupervised clustering of the CyTOF data (see ‘CyTOF’ below).

#### EBOV serum viral load by RT-qPCR

70 μL of sample inactivated by TRIzol LS was added to 280 μL of Buffer AVL (Qiagen, Hilden, Germany) with carrier RNA (Bennett et al. in submission). Samples were then extracted using the QIAamp Viral RNA Mini Kit (Qiagen) in accordance with the manufacturer’s instructions, eluted in 70 μL of Buffer AVE, aliquoted, and frozen. Viral load was determined using the BEI Resources Critical Reagents Program experimental EZ1 RT-qPCR kit assay in accordance with the manufacturer’s instructions.

#### PBMC purification

We centrifuged whole blood in K_3_EDTA tubes at 1800 x *g* for 10 min at room temperature, removed EDTA plasma, and added phosphate buffered saline (PBS, Thermo Fisher Scientific, Waltham, MA, USA) to the pelleted cells to double the original whole blood volume. We gently poured the PBS-blood cell mixture into an Accuspin tube containing Histopaque (Sigma-Aldrich, St. Louis, MO, USA) and centrifuged at 1000 x *g* for 10 min at room temperature with the brake set to 1. Following centrifugation, we removed the top, clear supernatant layer to within 0.5 cm of the cloudy white layer containing PBMCs.

We transferred the cloudy PBMC layer to a clean 15 mL conical tube and increased the volume to 10 mL using PBS supplemented with 2% heat-inactivated fetal bovine serum (PBS/2%HI-FBS) and mixed by inversion. We then centrifuged at 300 x *g* for 10 min at 4 °C with the brake set to 1. Following centrifugation, we removed the supernatant, resuspended the cell pellet with PBS/2%HI-FBS to a final volume of 10 mL, and mixed using gentle raking to wash the cells. We repeated the wash step 2 more times with the centrifuge set to 200 x *g* for 10 min at 4 °C with the brake set to 1. We then resuspended the cell pellet in 9.5mL PBS/2%HI-FBS for counting using the Countess Cell Counting system (Thermo Fisher Scientific). We aliquoted 0.5 mL for Seq-Well, and used the remaining 9 mL volume for CyTOF.

#### Seq-Well

We performed Seq-Well as described previously (Gierahn et al., 2017), with the S^3^ protocol (Hughes et al., 2019) and some controls and modifications to adhere to the BSL-4 environment.

As an experimental negative control to test our statistical model, we spiked Madin-Darby canine kidney (MDCK) cells, constituting ∼5% of the total sample, into a subset of PBMC samples (**Table S1**) from EVD NHPs immediately before scRNA-Seq. As MDCKs were not exposed to EBOV, viral reads in these transcriptomes should be due to ambient RNA contamination (Russell et al., 2019).

After loading and sealing beads and cells in Seq-Well arrays, we placed them in a −80 °C freezer until further processing – this step was required due to time constraints in the BSL-4. Later, we removed sealed Seq-Well arrays from the −80 °C freezer, placed them in 4-well dishes, and allowed them to equilibrate to room temperature for at least 30 min. We then covered arrays in 5 mL Seq-Well Lysis Buffer per protocol.

We performed RNA hybridization and RT as specified in the protocol (Hughes et al., 2019). After RT, we collected beads by centrifugation at 1000 x *g* for 1 min at room temperature. We resuspended beads with GeneXpert Lysis Buffer (Cepheid, Sunnyvale, CA, USA) for inactivation, which was required prior to removal from the BSL-4 laboratory according to standard operating procedures. After removal, we washed beads thrice with TE buffer containing 0.01% Tween 20 and shipped at 4 °C for further library construction and sequencing, which was performed with the S^3^ protocol (Hughes et al., 2019). We sequenced all libraries on either NextSeq 550 High Output or NovaSeq 6000 S2 flowcells (Illumina, San Diego, CA, USA), with 20 cycles for Read 1 (cell barcode and unique molecular index [UMI]) and 88 cycles for Read 2 (cDNA of interest). In some cases, we merged fastq files from multiple sequencing runs for increased coverage.

#### Depletion of abundant sequences by hybridization (DASH) of select Seq-Well libraries

We observed long concatemers of the common scRNA-Seq adaptor sequence (SeqB, 5’-AAGCAGTGGTATCAACGCAGAGTAC-3’) at high frequency in some Seq-Well libraries, likely owing to low RNA input and the challenging environment of processing samples in the BSL-4 suite. These concatemers disrupted Illumina sequencing runs because the Read 1 sequencing primer annealed to multiple SeqB sequences on a single template, allowing multiple sequencing-by-synthesis reactions simultaneously. We therefore devised a strategy to remove SeqB concatemers using depletion of abundant sequences by hybridization (DASH) (Gu et al., 2016), a CRISPR-based method to degrade target DNA sequences prior to sequencing. SeqB lacked a ‘NGG’ protospacer adjacent motif (PAM) for the common *S. pyogenes* Cas9 (SpyCas9) for which DASH was originally described; therefore, we modified DASH to use *S. aureus* Cas9 (SauCas9) (Ran et al., 2015). Moreover, in contrast to SpyCas9, SauCas9 is a multi-turnover enzyme (Yourik et al., 2019), suggesting that SauCas9 would have higher cleavage efficiency, which was important since SeqB was present in multiple copies within a concatemer.

First, we designed a guide RNA (gRNA) to target SeqB. Based on the position of the SauCas9 PAM and the length of SeqB, only 17 nucleotides of the gRNA protospacer could anneal to SeqB. Because gRNA length is critical to SauCas9 cleavage efficiency (Friedland et al., 2015; Ran et al., 2015), we prepended 4 random bases as a 5’ overhang (**Key Resources Table**). We *in vitro* transcribed this gRNA using the MEGAshortscript T7 Transcription Kit (Thermo Fisher Scientific), purified it using the RNA Clean & Concentrate Kit (Zymo Research, Irvine, CA, USA), and verified the correct RNA length on a 15% TBE-urea gel (Bio-Rad Laboratories, Hercules, CA, USA).

We performed SauCas9 DASH according to reaction conditions laid out for *in vitro* SauCas9 cleavage assays (Yourik et al., 2019). We incubated 10 pmol gRNA with 5 pmol SauCas9 (New England Biolabs [NEB], Ipswich, MA, USA) at 25 °C for 10 min, and then added up to 5 fmol DNA (2000:1000:1 RNA:protein:DNA ratio) and NEBuffer 3.1 (NEB) to 1X. We incubated this reaction at 37 °C for 2 hours (h), and quenched by adding EDTA to 50 µM, SDS to 1%, and 4 total U of Proteinase K (NEB) at room temp for 10 min. We removed degraded concatemers with two consecutive 0.8X SPRI purifications using Ampure XP beads (Beckman Coulter, Brea, CA, USA), eluted, and performed 6–9 cycles of PCR with the NEBNext Ultra II Q5 Master Mix (NEB) using Illumina P7 and the Seq-Well P5-TSO hybrid primer (Gierahn et al., 2017).

#### CyTOF

We added 1 mL of 16% paraformaldehyde (PFA, Electron Microscopy Sciences, Hatfield, PA, USA) to 9 mL PBMCs to fix the cells. We incubated samples at room temperature for 10 min followed by a final centrifugation at 600 x *g* for 5 min at 4 °C with the brake set to 9. We removed the supernatant, added 1 mL of PBS/5%HI-FBS for every 3 x 10^6^ cells (*e.g.*, 2 mL for a sample containing 6 x 10^6^ cells), aliquoted into 1 mL aliquots in cryovials, and stored in a −80 °C freezer.

We equilibrated fixed PBMCs to come to room temperature before transferring approximately 2 x 10^6^ cells per sample into 1.2 mL cluster tubes in a 96-tube rack. We barcoded samples and multiplexed them into 6 batches of 16 samples using a previously described 16-plex palladium-based mass-tag cell barcoding scheme (Zunder et al., 2015). We aspirated pelleted barcoded cells to a volume of 50 µL and incubated them with 15 µL of Human TruStain FcX (Biolegend, San Diego, CA) for 10 minutes. We stained cells for 30 min with 175 µL of a reconstituted lyophilized cocktail of metal-tagged cell-surface antibodies described previously (Bjornson-Hooper et al., 2019a, 2019b) supplemented with two additional antibody channels (**Table S7**). We washed and permeabilized surface-stained cells with methanol before aspirating down to 50 µL and staining for 60 min with 190 µL of reconstituted lyophilized intracellular staining antibody cocktail (Bjornson-Hooper et al., 2019a, 2019b) (**Table S7**). We washed and resuspended fully-stained cells in a volume of 750 µL.

Following staining, we inactivated samples by adding 250 μL of 16% PFA to 750 μL of each sample, for a final concentration of 4% PFA, and incubated at 4 °C overnight. The next day, we centrifuged samples 600 x *g* for 5 min at 4°C and aspirated down to 100 μL. We resuspended samples in 1 mL 4% PFA in PBS and transferred to a clean 2 mL cryovial. We then removed samples from the BSL-4 using a dunk tank and froze at −80 °C within 30 min of PFA addition.

Following inactivation, we thawed and processed inactivated samples within a BSL-2 lab for iridium intercalation, then mixed with 1xEQ beads (Fluidigm, South San Francisco, CA, USA) and run on a CyTOF Helios (Fluidigm) instrument using a Super-Sampler introduction system (Victorian Airship & Scientific Apparatus LLC, Alamo, CA, USA).

FCS data files were normalized across all runs using the data normalization software (Finck et al., 2013) and debarcoded using the single-cell debarcoder tool (Zunder et al., 2015) as previously described. Data were uploaded and analyzed using CellEngine software (https://cellengine.com, Primity Bio, Fremont, CA, US), and a gating strategy was applied to identify cell populations using canonical markers. Frequencies for each population were determined as a function of total CD66-CD45+ cell events and reported marker intensities are expressed as medians. Also see ‘CyTOF data preprocessing, clustering, and dimensionality reduction’ section below for details of unsupervised analysis of CyTOF data.

#### *Ex vivo* inoculation of PBMCs

PBMCs were isolated from healthy NHPs as previously described. We diluted cells to 3.3 x 10^6^ cells/mL RPMI/10%HI-FBS and transferred 900 µL each to 2 mL external thread cryogenic vials (Corning, Corning, NY) for each experimental condition. We inoculated cells with 100 µL live virus (EBOV/Kikwit, the same stock used for *in vivo* NHP inoculation) for a final MOI of 0.1 PFU/cell, an equivalent dose of irradiated virus (EBOV/Kikwit treated with 5 mRads gamma irradiation), or media only, and incubated for 4 or 24 hours with slow rocking prior to Seq-Well processing.

#### Single-cell RNA-Seq raw data processing

Raw sequencing files were demultiplexed and converted to fastq using bcl2fastq version 2.20. Reads were then trimmed, aligned to a reference transcriptome, and parsed into a digital gene expression matrix using the previously published Dropseq-tools pipeline (Macosko et al., 2015) version 2.0. We used a Snakemake wrapper around Dropseq-tools that is available in the open source Github repository https://github.com/Hoohm/dropSeqPipe. In brief, we trimmed adapter sequences using Cutadapt (Martin, 2011) version 1.16, performed spliced alignment of trimmed reads using STAR aligner (Dobin et al., 2013) version 2.6.1b, identified core barcodes using the whitelist function in umi_tools (Smith et al.) version 0.5.5, and used Dropseq-tools to correct barcodes and extract digital count matrices. A frozen version of the pipeline used to process the Seq-Well data is available at the Github repository https://github.com/dylkot/dropSeqPipe-dak.

Sequencing reads were aligned to a hybrid genome/genebuild of *Macaca mulatta* (genome assembly Mmul_8.01, Ensembl gene build 92) and EBOV/Kikwit (Genbank accession KU182905.1).

#### Single-cell RNA-Seq data preprocessing, clustering, dimensionality reduction, and smoothing

The scRNA-Seq data was preprocessed, clustered, and visualized using Scanpy (Wolf et al., 2018). We removed cells with <300 genes detected, >10% of their UMIs derived from mitochondrial genes, or >95% of UMIs mapped to non-genic regions. We excluded ribosomal genes, genes correlated with the percentage of UMIs assigned to mitochondrial genes (Pearson R > 0.1), and *HBB* as these were largely driven by the technical covariate of whether cells had loaded into Seq-Well arrays fresh or had undergone a freeze-thaw cycle with cryoprotectant. We also excluded EBOV genes and cell-cycle genes (defined by correlation with *TOP2A,* Pearson R > 0.1) prior to clustering so that these signals would not influence identification of cell types.

We performed multiple iterations of clustering to detect and exclude doublets and to identify distinct cell populations at multiple levels of granularity. In each iteration, we filtered genes detected in <10 of the cells being clustered. We then transformed raw UMI counts by normalizing the sum of counts of each cell to 10,000 (TP10K), adding 1 to each expression value, and taking the natural logarithm. In each clustering iteration, we identified and subsetted the data to highly variable genes using the highly_variable_genes function in Scanpy (Satija et al., 2015) with the default parameters. We Z-normalized each gene and set transformed values exceeding 10 to 10. Z-normalized data was used as input to principal component analysis. We determined the number of principal components to use for downstream analysis by identifying an elbow on the Skree plot of the eigenvalues associated with each principal component. For the *in vivo* Seq-Well data, we used the Harmony algorithm (Korsunsky et al., 2019) to remove variation due to whether a PBMC sample had been processed fresh, or following a freeze-thaw. No Harmony adjustment was used for the *ex vivo* EBOV dataset. The Harmony-adjusted or raw principal components were then used to construct a nearest neighbor graph with the number of neighbors set to the maximum of 30 or 0.001 x the number of cells. Lastly, we clustered cells using the Leiden community detection algorithm (Traag et al., 2019).

We annotated broad PBMC clusters in the *in vivo* and *ex vivo* EBOV datasets based on the following marker genes: CD8+ T-cells (*CD3D*, *GZMB*, *GNLY*), CD4+ T-cells (*CD3D*, *IL7R*), B-cells (*MS4A1*, *IGHM*), Monocytes (*CFD*, *LYZ*), cDCs (*FLT3*, *IRF8*), pDCs (*IRF8*, *GZMB*), Neutrophils (*CD177*, *LCN2*), Platelets (*PF4*, *CAVIN2*), Plasmablasts (*MZB1*, *IGHM*, *IGHA*), and spike-in control MDCK cells (*COL5A2*, *SLC20A1*).

For the *in vivo* EBOV dataset, the first clustering iteration was used to identify and filter a cluster of multiplets (expressing high levels of B-cell, T-cell, Neutrophil, and Monocyte genes) and MDCK control cells. A second clustering iteration was run on the filtered data to identify broad cell type clusters of T/NK-cells, B-cells, and myeloid cells (Monocytes, cDCs, pDCs, and neutrophils). Sub-clustering was performed on each broad cell type in two iterations, the first to identify remaining doublets to exclude, and the second to cluster cells into final cell-types and sub-types based on annotation of marker genes (Figure S4).

An analogous sequence of clustering iterations was used for the *ex vivo* EBOV dataset. However, as there were no MDCK cells spiked in, we proceeded straight to sub-clustering of the T/NK, B, Monocyte/DC, and multiplet populations from the initial clustering iteration.

Doublets and multiplets identified during any clustering iteration were excluded. Then visualization in 2 dimensions was accomplished by computing the nearest neighbor graph with 0.001 x the number of cells as nearest neighbors, followed by Uniform Manifold Approximation and Projection (Becht et al., 2018).

Gene expression values were smoothed to facilitate direct visualization of *CD14*, *FCGR3* (which codes for CD16), and *MKI67* (which codes for Ki67) by running MAGIC (van Dijk et al., 2018) on log TP10K expression values with the ‘cosine’ distance metric and 3 diffusion steps.

#### Differential expression testing

We performed all differential expression tests using MAST (Finak et al., 2015) on log(TP10K + 1) normalized data. For all differential expression tests, we included (1) the percentage of mitochondrial reads and (2) the number of genes detected in a cell as covariates. For tests in the *in vivo* dataset, we additionally included a binary indicator covariate of whether the cell was derived from a sample that had been processed fresh, or had undergone freeze-thaw. For tests in the *ex vivo* dataset, we additionally included a binary indicator covariate of which NHP donor the sample was derived from.

For viral load comparisons, we used the log10 viral load as a continuous exogenous variable and only considered cells with ≥1 viral read. For all other comparisons, we used a binary exogenous variable indicating the reference and the query group. “Viral load” and “bystander vs. EBOV infected cells” in the *in vivo* dataset were conducted only considering cells from DPI 5–8. “Viral load” comparisons in the *ex vivo* dataset only considered cells from the 24 HPI timepoint.

Differential expression *p*-values were corrected for multiple hypothesis testing using the method of Benjamini and Hochberg (Benjamini and Hochberg, 1995).

#### Identifying differential gene expression modules

We clustered the log fold-change profiles of 1,437 differentially expressed genes (rows of Figure 3A) considering B, CD4+ T, CD8+ T, NK, and monocyte populations at the three EVD stages. Prior to clustering, insignificant values (*p* > 0.2) were first set to 0 and genes were normalized to unit L2 norm. We performed KMeans clustering with K=11 and the default parameters in Scikit-learn (Pedregosa et al., 2011) version 0.22.2.post1. Varying K above and below 11 led to highly consistent results with splitting or merging individual clusters at the margin, and we picked K=11 as the lowest value that yielded the “B down” module.

#### Detection of EBOV infected cells

Our infection detection model assumes that EBOV reads assigned to a cell are either due to true viral RNAs inside that cell, or due to ‘ambient’ extracellular EBOV RNA in the sample that, by chance, are captured in a well of the Seq-Well array along with the cell. Our null model for how many EBOV reads would be expected in a cell by chance therefore depends on two main parameters which we estimate from the data: (1) what proportion of ambient RNAs in a sample are due to EBOV, and (2) what proportion of a cell’s expression profile is due to ambient RNA. Our method therefore precedes through the following steps which we describe below:

1. Estimate an ambient RNA profile for each Seq-Well array
2. Estimate the proportion of each cell’s transcripts that are due to ambient RNA
3. Determine if there are more EBOV transcripts in each cell than would be expected based on the proportion of EBOV in the ambient RNA and the cell’s estimated level of ambient RNA contamination.

##### 1. Estimate an ambient RNA profile for each Seq-Well array

We assume that cell barcodes with few transcripts detected correspond to wells in the Seq-Well array that lack a cell, and therefore, that any transcripts assigned to those cell barcodes derive from ambient RNA. We consider all cell barcodes with fewer than 50 transcripts detected to be empty wells and compute an ambient RNA profile of the proportion of transcripts from these barcodes assigned to each gene. This is analogous to the approach used in several ambient RNA correction approaches such as (Fleming et al., 2019).

We denote the number of genes in the dataset as *G* and define the ambient RNA profile for a given Seq-Well array, *a*, as a *G*-dimensional vector 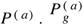 denotes the estimated proportion of ambient RNAs that are assigned to gene *g* based on transcripts from all cell barcodes with fewer than 50 UMIs. Since 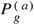 is a proportion, the following constraints hold:

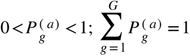

##### 2. Estimate the proportion of each cell’s transcripts that are due to ambient RNA

We adapt the previously published Consensus Non-negative Matrix Factorization (cNMF) method (Kotliar et al., 2019) to estimate the ambient RNA contamination level of each cell. In brief, cNMF learns a user-specified number of gene expression programs (GEPs), each a non-negative *G*-dimensional vector representing the average expression profile of an individual cell type or cellular activity (e.g. cell-cycle or interferon response) that are present in the data. In addition, it learns a “Usage” matrix reflecting the % contribution of each GEP in each cell (*i.e.*, the % of each cell’s transcripts derived from each GEP). For the following notation, matrices are indicated in bold. Denoting the number of cells in the dataset as *C* and the user-specified number of GEPs as *K*, cNMF is given an input *C*x*G* matrix of transcript counts 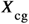 and returns 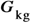 a non-negative *K*x*G* matrix of GEPs reflecting the relative contribution of gene *g* in GEP *k*, as well as a non-negative *C*x*K* matrix 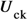 reflecting the percentage of transcripts in cell *c* that are due to GEP *k*.

We adapt this approach to return an updated, *C*x(*K+1*) dimensional usage matrix that includes an additional column reflecting the usage of the ambient RNA profile. We run cNMF as published to obtain the 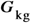 GEP matrix. We then append the ambient RNA profile for a given array as an additional row to 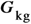 and normalize each row to sum to 1 like so:

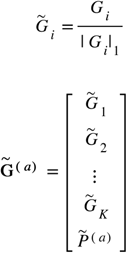

where 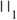 denotes the L1 norm, a tilde is used to denote an L1 normalized vector, and 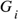 denotes the ith row of 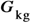. We then run a final iteration of NMF with the GEP matrix fixed to 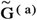. This jointly estimates a usage of the ambient RNA profile and the other GEPS using non-negative least squares. For a given cell *c* from an array *a*, this amounts to solving the following non-negative least squares optimization:

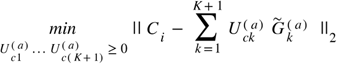

We combine these coefficients for all cells and programs into a single matrix and L1 normalize the usages to sum to 1:

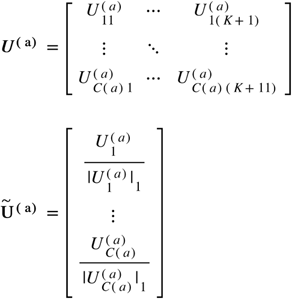

where 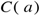 denotes the number of cells derived from array *a*. The “K+1”th column of 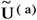 reflects the estimated proportion of transcripts due to ambient RNA in all of the cells from array *a.* We repeat this calculation for each array separately and denote the estimated contribution of ambient RNA for a cell *c* as 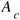.

##### 3. Determine if there are more EBOV transcripts in each cell than would be expected by chance

For each cell, we determine a threshold, 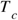, for the number of EBOV transcripts required to call that cell infected while keeping the false positive rate below *f*, a user specified threshold (*f =* 0.01 in all of our analyses). We first calculate 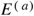, the proportion of ambient RNA transcripts in array *a* that are due to EBOV as follows:

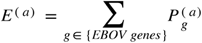

Then, for a cell with 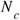 transcripts total; a given proportion, 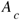, of its reads derived from ambient RNA; 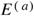, the proportion of ambient RNA contaminating reads expected to map to EBOV, we compute 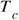 using binomial statistics as follows:

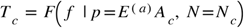

Where *F* is the inverse survival function of the Binomial distribution with event probability *p* and *N* trials. We identify cells with 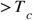 reads as infected.

#### Estimation of the infection receiver operator characteristic

We estimate sensitivity to call an infected cell with either 1% or 0.1% of its reads due to EBOV across a range of 60 false positive rate thresholds (*f*). We first randomly sampled 2,000 cells from the live EBOV treatment samples in the *ex vivo data,* or the non-baseline samples from the *in vivo* data, to serve as an empirical distribution for 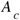 and 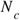. For each cell and specificity threshold *f*, we then calculate the probability of correctly calling a true positive cell as positive. We model the distribution of the number of EBOV reads in a cell as the convolution of 2 binomial distributions: 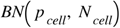 is the binomially distributed number of true cell-derived reads mapping to EBOV and 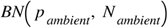 is the analogous distribution for ambient RNA-derived reads mapping to EBOV. 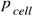 is 0.01 or 0.001 by assumption, and 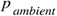 is estimated empirically for each cell as 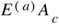. We calculate the expected values for 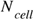 and 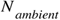 and round 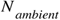 up to the nearest integer, as follows:

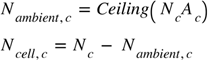

We then directly calculate the convolution of the two binomial distributions for each cell:

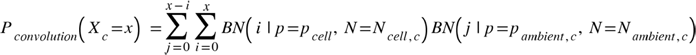

We then calculate the sensitivity for each cell as the probability that the convolution distribution is greater than the empirical threshold:

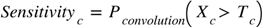

We plot the average sensitivity across the 2000 randomly sampled cells as a function of *f* for 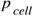 equal to either 0.01 or 0.001.

#### Gene set enrichment testing

We downloaded gene sets from the Molecular Signatures Database (Liberzon et al., 2011) version 6.2 for gene set enrichment testing. We considered all Hallmark or C2 gene sets containing greater than 10 genes that were present in our expression data. We tested expression modules for enrichment using Fisher’s exact test and corrected for multiple hypothesis testing using the method of Benjamini and Hochberg (Benjamini and Hochberg, 1995).

To test continuous expression profiles for gene set enrichment (Figure S6B), we used the rank-sum test comparing genes in the gene set to all genes not in the set.

#### Scoring cells for interferon response and macrophage differentiation

We identified 58 genes in the “Global up” module that were also included in one or more of the following gene sets from the molecular signatures database: HECKER_IFNB1_TARGETS, BROWNE_INTERFERON_RESPONSIVE_GENES, MOSERLE_IFNA_RESPONSE, HALLMARK_INTERFERON_ALPHA_RESPONSE, HALLMARK_INTERFERON_GAMMA_RESPONSE (**Table S3**). We then scored cells for the average expression of these genes using the score_genes function in Scanpy (Satija et al., 2015) with 58 control genes, as this was the number of genes in the ISG set, and otherwise default parameters.

We computed a macrophage score based on the set of 618 genes annotated as significantly up or down-regulated during *in vitro* monocyte-to-macrophage differentiation (Dong et al., 2013) (**Table S4)**. We computed each cell’s macrophage score as the dot-product of its expression profile for the 618 genes (in log TP10K) with the log fold-change reported for each gene in (Dong et al., 2013). This effectively weights genes by both the direction and magnitude of their change during *in vitro* macrophage differentiation.

#### Comparison of EVD monocyte subsets with human bone marrow and PBMC data

We obtained all of the human PBMC datasets produced using v3 or v3.1 chemistry from the 10X website (**Key Resources Table**), aggregated them together, and processed the resulting dataset using the same pipeline as the NHP Seq-Well data. Briefly, we first filtered out genes detected in fewer than 10 cells before converting to log TP10K and performed PCA as described above. Then, we used Harmony (Korsunsky et al., 2019) to integrate out variation due to the different samples of origin and used 30 nearest neighbors for Leiden community detection (Traag et al., 2019) and UMAP dimensionality reduction (McInnes et al., 2018) (Figure S7E). We did not perform any sub-clustering on this dataset.

We obtained Human Cell Atlas bone marrow data from the Human Cell Atlas data portal (Hay et al., 2018) and processed it according to the same pipeline as the NHP data with a few modifications. We filtered doublets prior to clustering by running Scrublet (Wolock et al., 2019) separately within each of 8 donor batches with an expected doublet rate parameter of 6%. We identified and excluded cell-cycle associated genes, as those with a Pearson correlation > 0.3 with *TOP2A*. We integrated data from the different donor batches using Harmony and used 30 nearest neighbors for Leiden community detection and UMAP dimensionality reduction. We performed 3 rounds of sub-clustering: First we clustered all of the cells to identify monocyte and dendritic lineage cells (Figure S7F). Second, we clustered just hematopoietic stem cells (HSCs) and monocyte/dendritic progenitor cells to identify doublets (as those falling into a cluster characterized by T-cell marker genes such as CD3D and CD3E). Finally, we re-clustered this set with the doublets excluded to identify monocyte lineage cells, plasmacytoid dendritic cells, and conventional dendritic cells (Figure S7G).

We confirmed our marker gene-based annotations of the myeloid cell populations by comparing these cells to the circulating human PBMC dataset. We identified the nearest neighbor of each bone marrow myeloid progenitor cell in the PBMC dataset based on Euclidean distance of TP10K-normalized cells, considering overdispersed genes identified in the PBMC dataset based on the V-score (baseline-corrected Fano factor) (Klein et al., 2015). We then visualized the nearest-PBMC assignment of the bone marrow myeloid cells on a UMAP embedding (Figure S7H).

Finally, we combined the monocytes and monocyte precursor cells from the human PBMC and bone marrow datasets into a single reference. We again normalized the data to log TP10K and computed UMAP embeddings following the same procedure as for the individual datasets (Figure S7I), using Harmony to remove variation due to donor sample. We then down-sampled this data so that there would be equivalent numbers of cells of each of the bone marrow and PBMC clusters (*i.e.*, 982 cells per cluster as that was the number of cells in the smallest cluster). We identified the nearest neighbor of each NHP monocyte in the down-sampled reference dataset, as described above and computed the percentage of NHP monocytes assigned to CD14+ or CD16+ clusters from either human bone marrow or PBMC (Figure 5F).

#### Logistic regression prediction of EBOV infection in vivo

We used Statsmodels (Seabold and Perktold, 2010) version 0.11.1 to fit a logistic regression predicting EBOV infection status among all monocytes from late EVD, based on the following features: macrophage score, MAGIC smoothed values of *CD14*, and *CD16* (*FCGR3*), and an interaction term for the product of the MAGIC smoothed values for *CD14* and *CD16*.

#### CyTOF data preprocessing, clustering, and dimensionality reduction

Our clustering and dimensionality analysis of the NHP CyTOF data was analogous to the Seq-Well pipeline with a few adaptations. We down-sampled a total of 1.1 million cells, consisting of 300,000 baseline cells and 100,000 cells from each DPI, selecting a uniform number of cells from each sample at a given DPI. We used the Arcsinh transformation of CyTOF raw intensity values divided by 5, which is standard in the field. We set a ceiling of transformed intensity values for each gene at the 99.999th percentile to reduce the effect of very rare outliers. We mean-centered the data but did not variance normalize prior to PCA. We then performed multiple clustering iterations of the data using the Leiden algorithm with the number of nearest neighbors set to the maximum of 30 or 0.01% of the number of cells in the dataset.

We annotated broad PBMC clusters in the CyTOF datasets based on the following marker proteins: CD8+ T-cells (CD3, CD8), CD4+ T-cells (CD3, CD4), NK cells (CD16, CD161), B-cells (CD19, IgM), Monocytes (CD11b, BDCA3, CD14, CD16, HLA-DR), cDCs (HLA-DR, CD11c, CD1c), pDCs (HLA-DR, CD4, CD123), Neutrophils (CD11b, CD66), Platelets (CD61, BDCA3), Plasmablasts (IgM high, but little or no CD19), Basophils (CD11b, CD123, low HLA-DR). There was also a cluster of cells characterized by high HLA-DR, Ki67, and CD38 which we annotate as “Unassigned APC”.

In the first clustering iteration, we identified and excluded clusters of doublets as those expressing markers of two or more broad cell-types. We also excluded a cluster characterized by high expression of CD3 but low expression of both CD4 and CD8 that we speculated was a technical artifact as it was nearly exclusive to a single CyTOF batch.

Next, to address batch effects between the different CyTOF runs, we grouped cells into the broad categories of Monocytes and DCs, CD4+ T, CD8+ T, NK, Neutrophils, B, Plasmablasts, and Platelets, and used COMBAT (Johnson et al., 2007) to adjust for batch effect separately within each broad category, using the default COMBAT parameters in the Scanpy implementation.

We then ran cNMF (Kotliar et al., 2019) to identify and regress out artifact signals in the data. We ran cNMF using all 42 markers as inputs and adapted the method to not perform any variance scaling prior to NMF; this is because CyTOF intensities of different markers are already expressed on a comparable scale, unlike genes in single-cell RNA-Seq data which can vary over different orders of magnitude. We also set a floor on the input data to be non-negative (because after COMBAT, a small percentage of the values were slightly below 0). We selected K=7 as the cNMF dimensionality as the solution stability fell off dramatically at higher values. One gene expression program (GEP) was characterized by high levels of the platelet markers CD61 and BDCA3. It was mostly enriched in the platelet cluster but was also elevated in a subset of cells of all major cell types, which suggests that it reflects an artifact of platelets sticking to cells. A second GEP was characterized by high levels of all of the intracellular markers and it was also distributed throughout cells of multiple clusters. We interpreted this GEP as a cell permeabilization artifact reflecting the relative accessibility of a cell’s intracellular proteins to CyTOF antibody staining. We regressed these 2 GEPs out of the data by subtracting the matrix (outer) product of the Usage and GEP matrices. We denote the number of cells as *C*, the number of genes as *G* (42 in our data), and the number of programs selected as K (7 in our data). The *C*x*G* input data matrix is denoted as, the *K*x*G* GEP matrix returned by cNMF as and the Cx*K* usage matrix as. Then the correction is as follows:

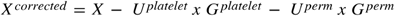

Where 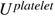 and 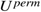are the *Cx*1 dimensional matrices representing the usage of the platelet and permeabilization GEPs 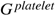 and and 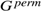are the 1x*G* dimensional matrices representing the spectra of the platelet and permeabilization GEPs, and multiplication is the outer product. We also filtered cells assigned to the platelet cluster in the initial clustering, since their predominant signal had been regressed out.

We then repeated clustering of the corrected data to identify broad cell types (Neutrophils, Monocytes, and DCs), followed by sub-clustering within each broad cluster to generate the sub-clusterings in Figure S4. The data was visualized using the UMAP algorithm as described for the scRNA-Seq data.

The human PBMC CyTOF data was processed with the same pipeline as the NHP data with a few modifications. The data were down-sampled to 280,000 cells total (20,000 per sample), batch correction was performed using Harmony, and 0.001 x the number of cells was used for K nearest neighbor graph construction. No cNMF or COMBAT adjustment steps were performed.

### Quantification and Statistical Analysis

Details of statistical testing, sample size, center, and dispersion can be found in the figure legends, the main text, and Materials and Methods.

### Additional Resources

This study did not generate new additional resources (website, forum, clinical trial).

### Key resources table

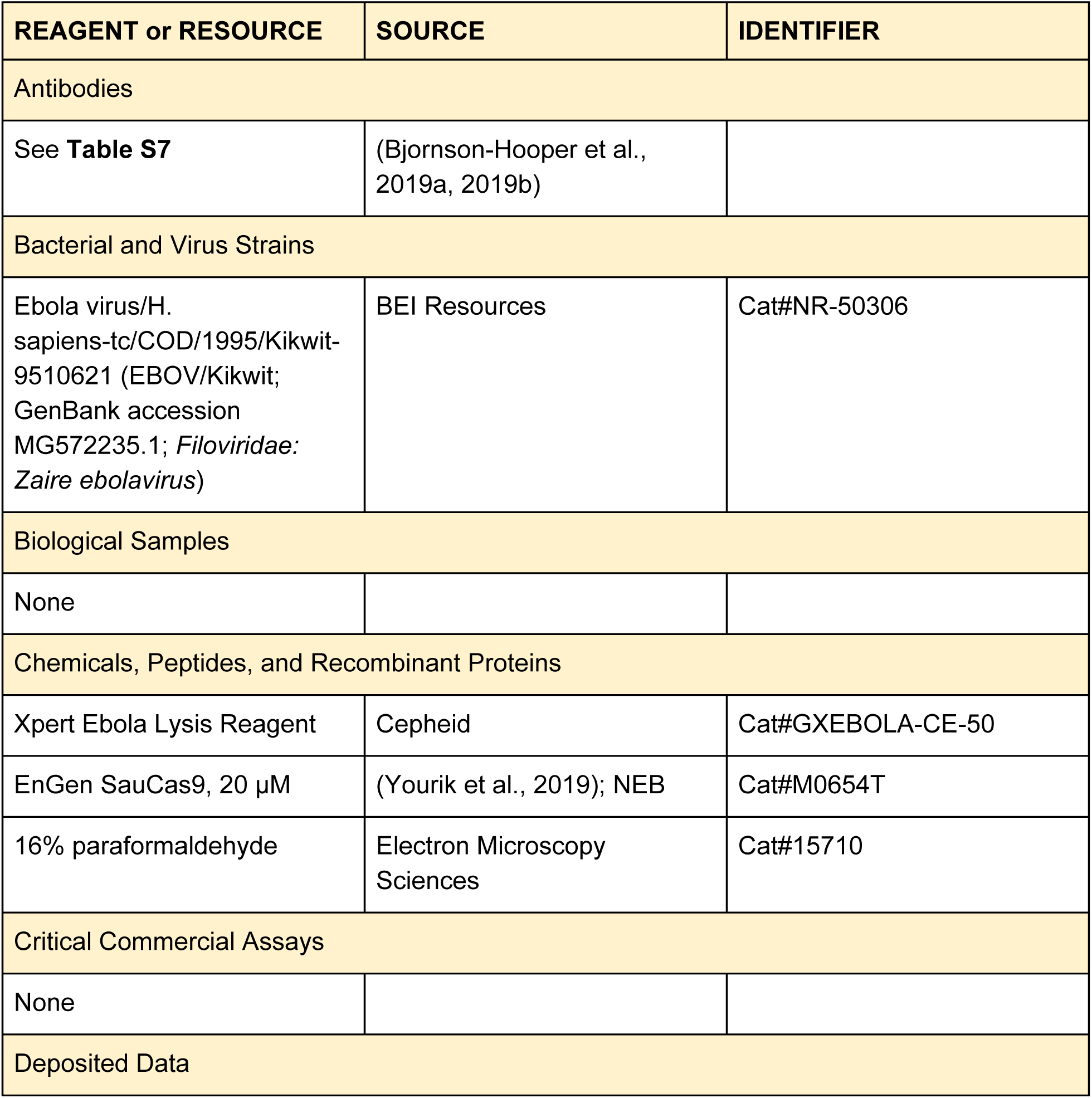
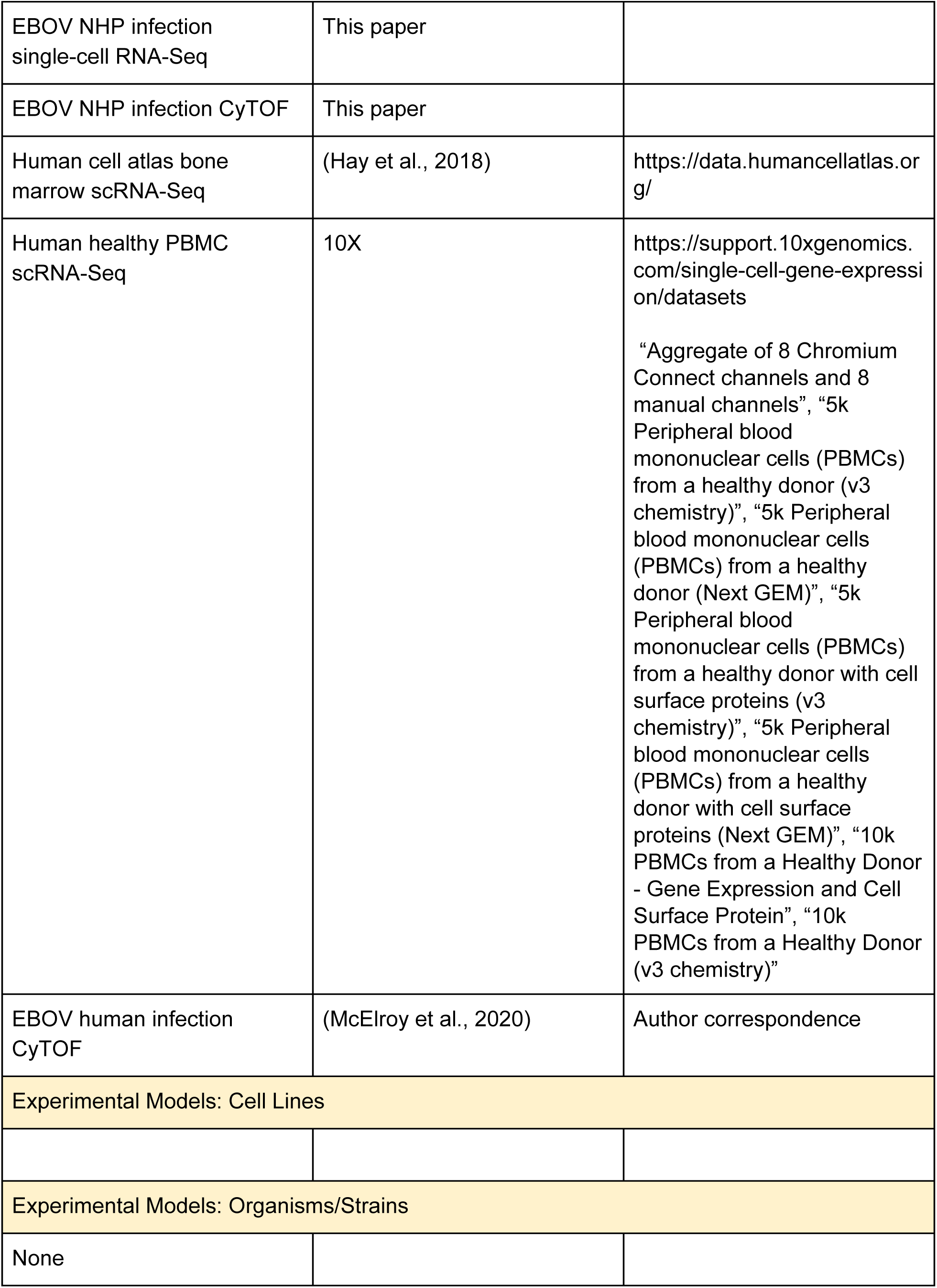
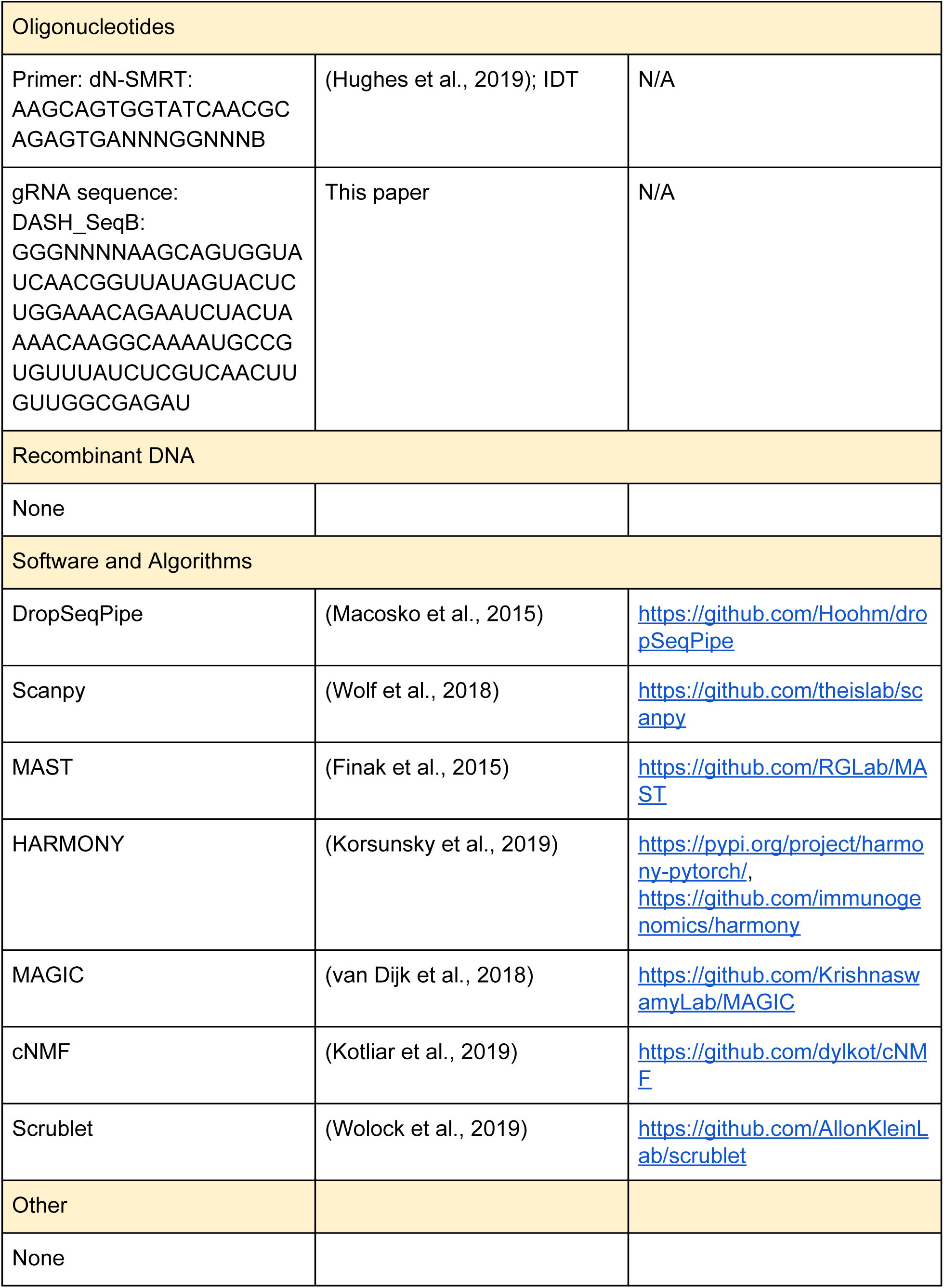

## Acknowledgements

We thank S. Schaffner, E. Normandin, K. Siddle, S. Reilly, S. Weingarten-Gabbay, C. Myhrvold, K. DeRuff, M. Rudy, N. Barkas, M. Babadi, M. Reyes, N. Hacohen, A. Regev, and E. Hodis, for helpful comments on this work. We thank S. Wolock for useful feedback and for providing scripts and processed data for investigating the human bone marrow samples. We thank D. Schwarz and New England Biolabs for generously providing SauCas9 and technical advice. We thank A. Matthews and M. Kemball for help in project management and administration. We thank S. Knemeyer and SciStories for illustrations.

This work is supported by the US Food and Drug Administration (FDA) contracts HHSF223201810172C and HHSF223201610018C, National Institute of Allergy and Infectious Diseases (NIAID) U19AI110818, and HHMI. This work was partially supported by NIAID Interagency agreement NOR15003-001-0000. The nonhuman primate work completed at the NIAID Integrated Research Facility was supported in part by the NIAID Division of Intramural Research and NIAID Division of Clinical Research and was performed under Battelle Memorial Institute Contract (No. HHSN272200700016I) and manuscript drafting was performed under Laulima Government Solutions, LLC. Contract (No. HHSN272201800013C). J.L. performed this work as an employee of Battelle. J.R.K., B. D.-K., R.A., and R.S.B. are current employees of Laulima Government Solutions, LLC. D.A.K. was supported by award Number T32GM007753 from the National Institute of General Medical Sciences (NIGMS). A.E.L. was supported by the National Science Foundation (NSF) under Grant No. DGE 1144152. M.M. was a Gilead Fellow of the Life Sciences Research Foundation. K.G.B was supported by a K01 (NIH-TW010853) and an ASTMH Shope fellowship. A.K.S. was supported by the Searle Scholars Program, the Beckman Young Investigator Program, a Sloan Fellowship in Chemistry, NIH 5U24AI118672, and the Bill and Melinda Gates Foundation. The CyTOF facility at the trans-NIH Center for Human Immunology is supported by funding from the Intramural Research Program of the NIH. The authors are solely responsible for the content of this paper, which does not necessarily represent the official views of the US Department of Health and Human Services (HHS), the NIH, the NIGMS, the FDA, or the institutions and companies affiliated with the authors.

## Author Contributions

Conceptualization, D.K., A.E.L., T.K.H., G.P.N., K.G.B., L.E.H., D.R.M., A.K.S., P.C.S., R.S.B.; Methodology, D.K., A.E.L., J.L., T.K.H., N.M.K., M.H.W., D.R.M., R.S.B.; Software, D.K.; Validation, S.S.R.; Formal Analysis, D.K., A.E.L., S.S.R., H.C., M.M., D.R.M.; Investigation, J.L., N.M.K., H.C., J.R.K., B.D.-K., Z.B.B., N.M., B.A.S., M.R.B., G.C.A., R.A., K.G.B., D.R.M., R.S.B.; Resources, T.K.H., M.H.W., Z.B.B., N.T.; Data Curation, D.K., A.E.L., D.R.M., R.S.B.; Writing – Original Draft, D.K., A.E.L.; Writing – Review & Editing, D.K., A.E.L., J.L., T.K.H., N.M.K., S.S.R., M.H.W., H.C., J.R.K., B.D.-K., Z.B.B., N.M., B.A.S., N.T., M.R.B., G.C.A., R.A., J.L.R., M.M., G.P.N., K.G.B., L.E.H., D.R.M., A.K.S., P.C.S., R.S.B.; Visualization, D.K., A.E.L., S.S.R., H.C., D.R.M.; Supervision, J.L.R., M.M., G.P.N., K.G.B., L.E.H., D.R.M., A.K.S., P.C.S., R.S.B.; Project Administration, D.K., A.E.L., J.L., N.T., K.G.B., R.S.B.; Funding Acquisition, D.K., A.E.L., M.M., G.P.N., K.G.B., L.E.H., D.R.M., P.C.S.

## Supplementary Figures

**Figure S1.**
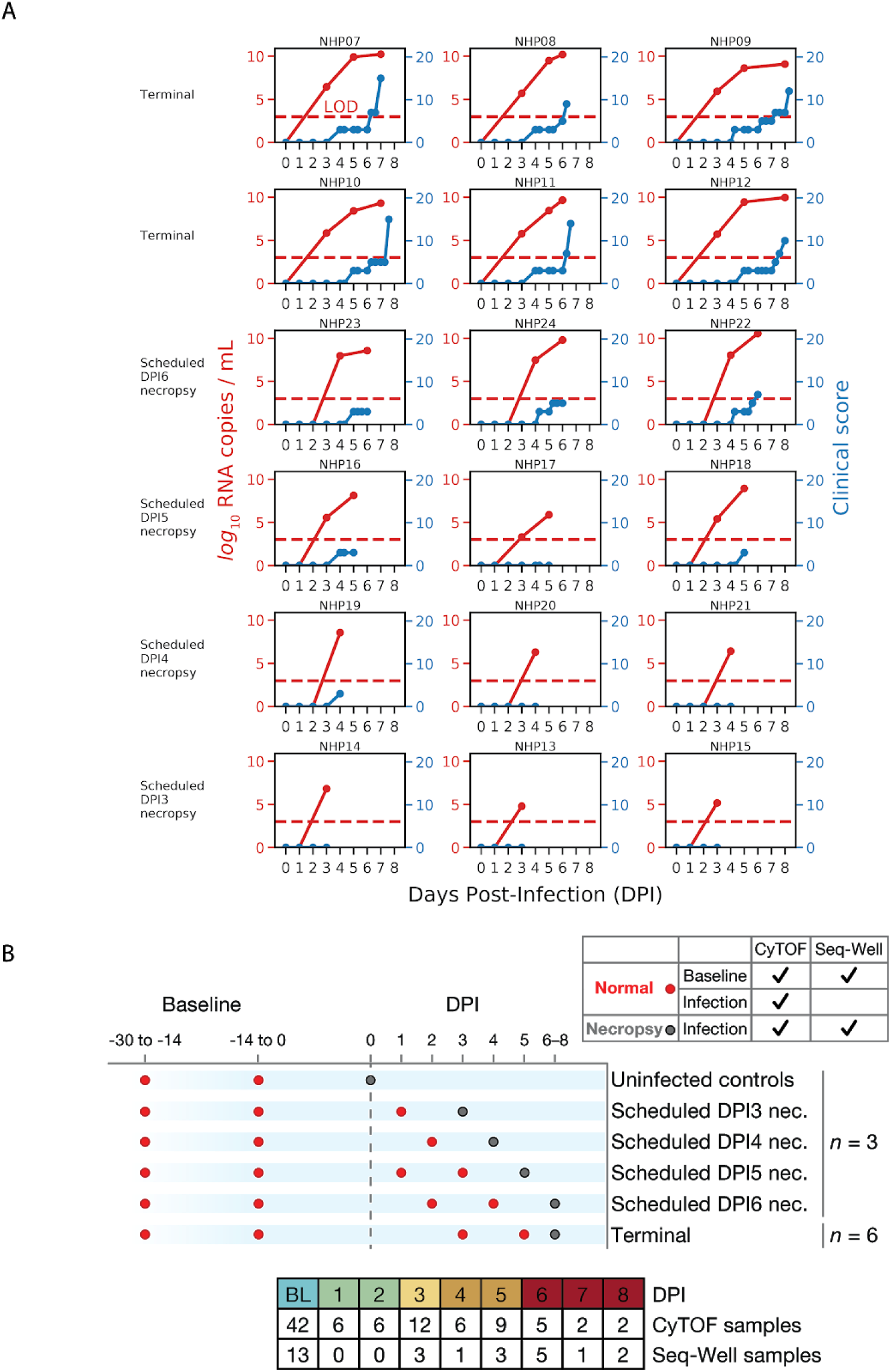
Blood sampling overview and clinical time course per animal replicate. Related to Figure 1. (**A**) Each panel represents the time course of log viral load (left axis, red) and clinical score (right axis, blue) for a specific NHP. Panels are organized into rows based on the cohorts. (**B**) Overview of study cohorts and blood draw timelines. Animals were grouped into cohorts with pre-scheduled necropsy times (at baseline, or day post infection [DPI] 3, 4, 5, 6 - *n* = 3 each), or allowed to progress until clinical score exceeded 10 (terminal), predetermined euthanasia criteria. Dots indicate scheduled blood draws for each cohort with red denoting an intermediate (non-necropsy) draw, and gray indicating a draw that coincided with euthanasia and necropsy. Necropsy and baseline normal draws were used for Seq-Well and CyTOF, while intermediate post-infection draws were available only for CyTOF.

**Figure S2.**
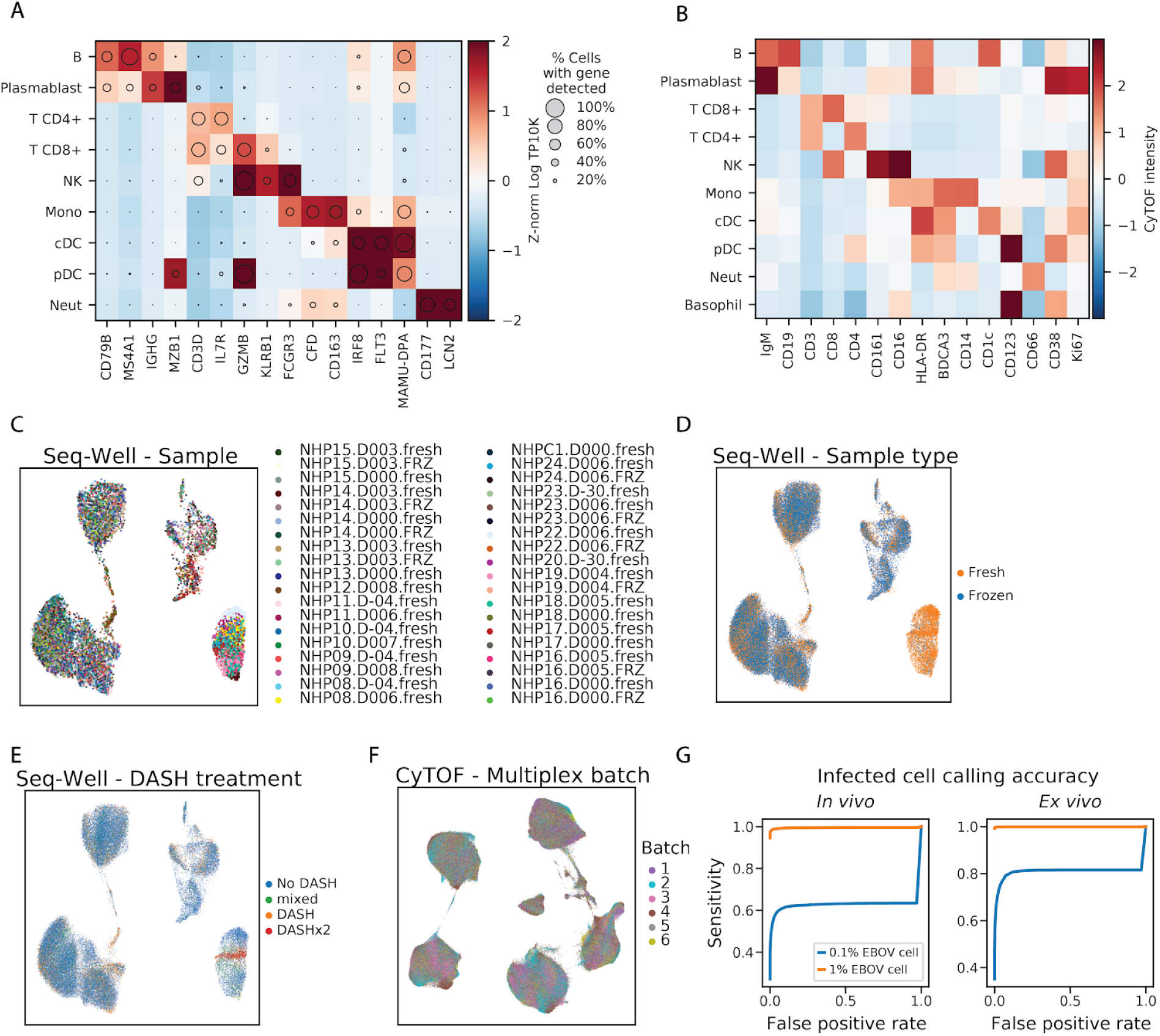
Cell type markers for Seq-Well and CyTOF clusters. Related to Figure 2. (**A**) Expression profiles of cell-type marker genes (columns) for cell-type clusters (rows) based on the *in vivo* Seq-Well data. Circle area represents the percentage of cells in each group in which the gene was detected, and color denotes the average expression level (natural log TP10K). (**B**) Average expression (Z-normalized CyTOF intensity) profiles of cell type marker genes (columns), for cell type clusters (rows), based on the CyTOF data. (**C**) Uniform Manifold Approximation and Projection (UMAP) embedding of post-integration Seq-Well data, colored by the sample source (NHP, DPI, and whether the sample was loaded for Seq-Well without any freezing [.fresh] or was frozen with cryoprotectant and thawed prior to Seq-Well [.FRZ]). A maximum of 500 cells per sample is plotted to increase representation across samples. (**D**) UMAP embedding of Seq-Well data, colored by whether cells were processed fresh (orange) or after freeze/thaw (blue) prior to Seq-Well. (**E**) UMAP embedding of Seq-Well data, colored by depletion of abundant sequences by hybridization (DASH) treatment. We developed a DASH-based method to remove a PCR adaptor artifact from some Seq-Well sequencing libraries (see Materials and Methods), and performed this 0 times (No DASH, blue), 1 time (DASH, orange), or 2 times sequentially (DASHx2, red). For a few samples, we sequenced ‘No DASH’ and ‘DASH’ libraries and merged the reads (mixed, green). (**F**) UMAP embedding of batch-corrected CyTOF data, colored by the multiplex batch in which it was pooled, and analyzed by CyTOF. (**G**) Receiver operating characteristic curves for identifying EBOV-infected cells. Estimates of sensitivity to detect an infected cell at various false positive rate thresholds *in vivo* (left) and *ex vivo* (right). Curves are estimated separately for a hypothetical viral load of 0.1% (blue line) and 1% (orange line).

**Figure S3.**
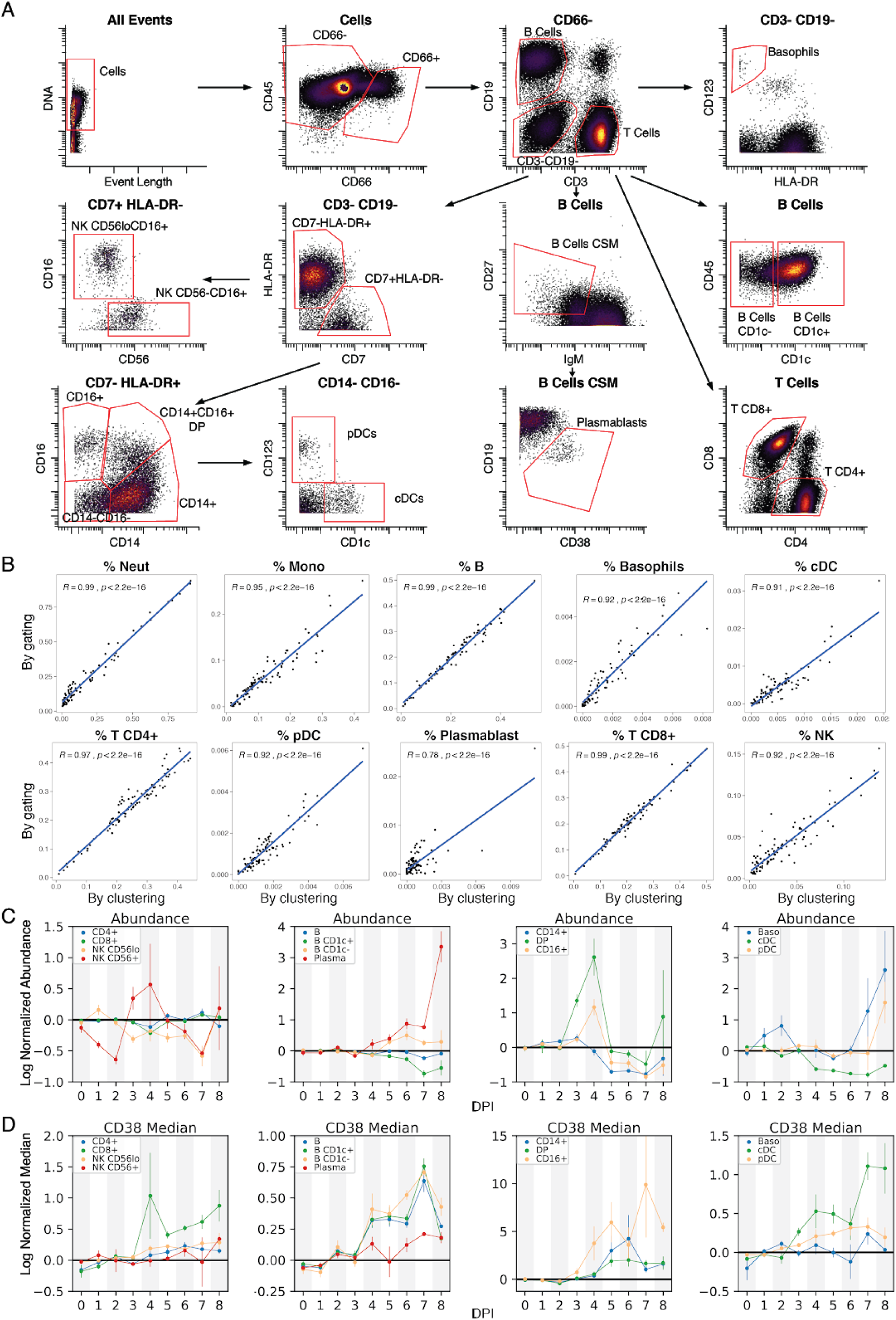
Comparison of unsupervised clustering and manual gating of CyTOF data. Related to Figure 2. (**A**) Gating strategy used to define cell populations using canonical markers. Axes indicate CyTOF ArcSinh-scaled marker intensities. (**B**) Pearson correlation between relative cell-type abundance determined by unsupervised clustering (*x*-axis) and by manual gating (*y*-axis). Each marker represents a single PBMC sample. (**C**) Abundances of indicated manually gated populations, normalized for each animal to pre-challenge timepoints. *x*-axis shows day post infection (DPI), lines denote mean ± 1 standard error of the mean (SEM). (**D**) Median CD38 expression on manually gated populations shown in (C), medians are normalized to each NHP’s average baseline expression. *x*-axis shows DPI, lines denote mean ± 1 SEM.

**Figure S4.**
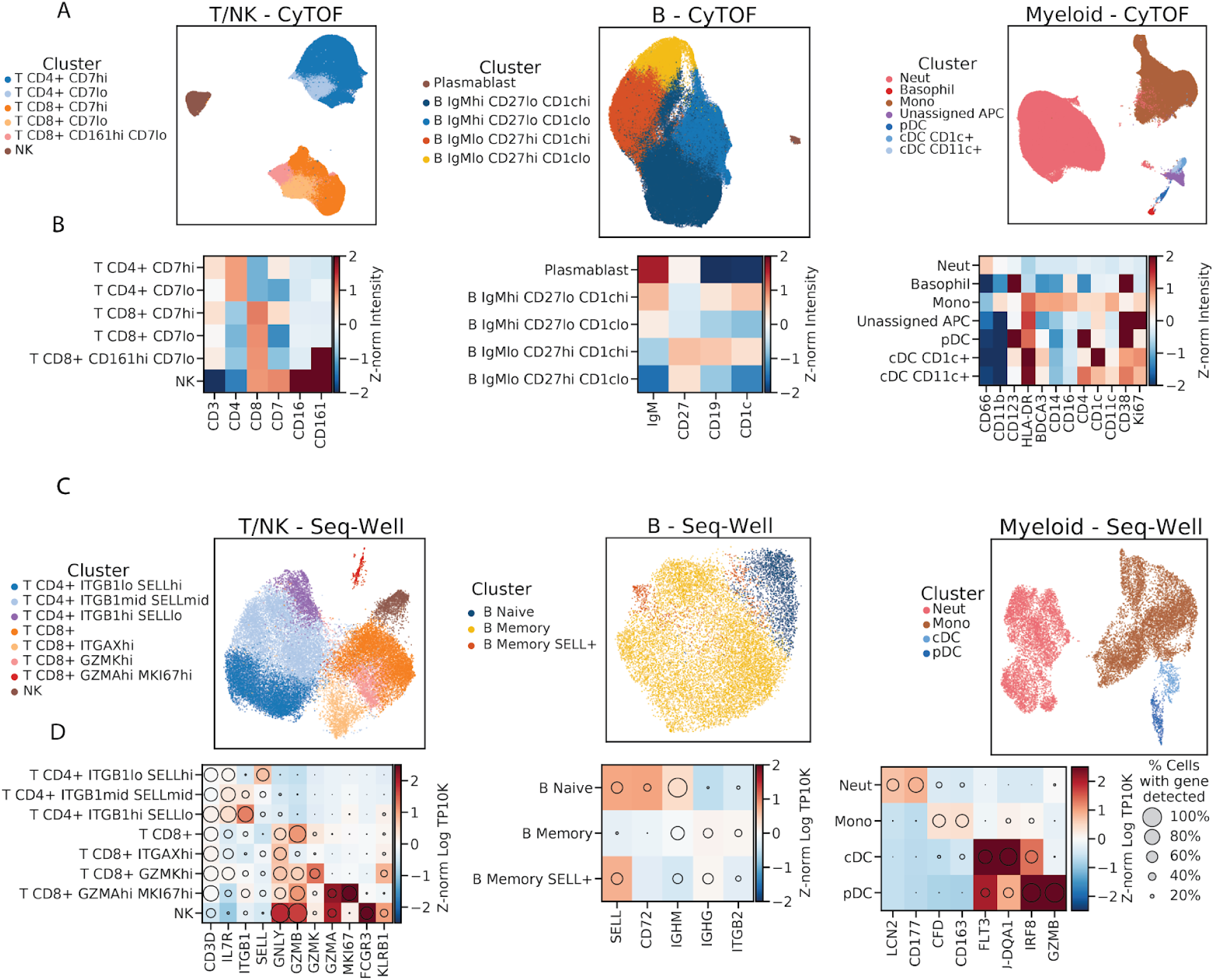
Identifying cell subtypes by subclustering. Related to Figure 2. (**A**) UMAP embedding of broad cell-type clusters in the CyTOF data, colored by sub-cluster assignment (Neut - neutrophil, Mono - monocyte). **(B)** Average expression (Z-normalized CyTOF intensity) profiles of sub-clusters for marker genes based on CyTOF data. **(C)** UMAP embedding of broad cell-type clusters in the Seq-Well data, colored by sub-cluster assignment. **(D)** Expression profiles of sub-clusters for marker genes based on Seq-Well data. Circle area represents the percentage of cells in which the gene was detected, and color denotes the average expression level (Z-normalized natural log TP10K).

**Figure S5.**
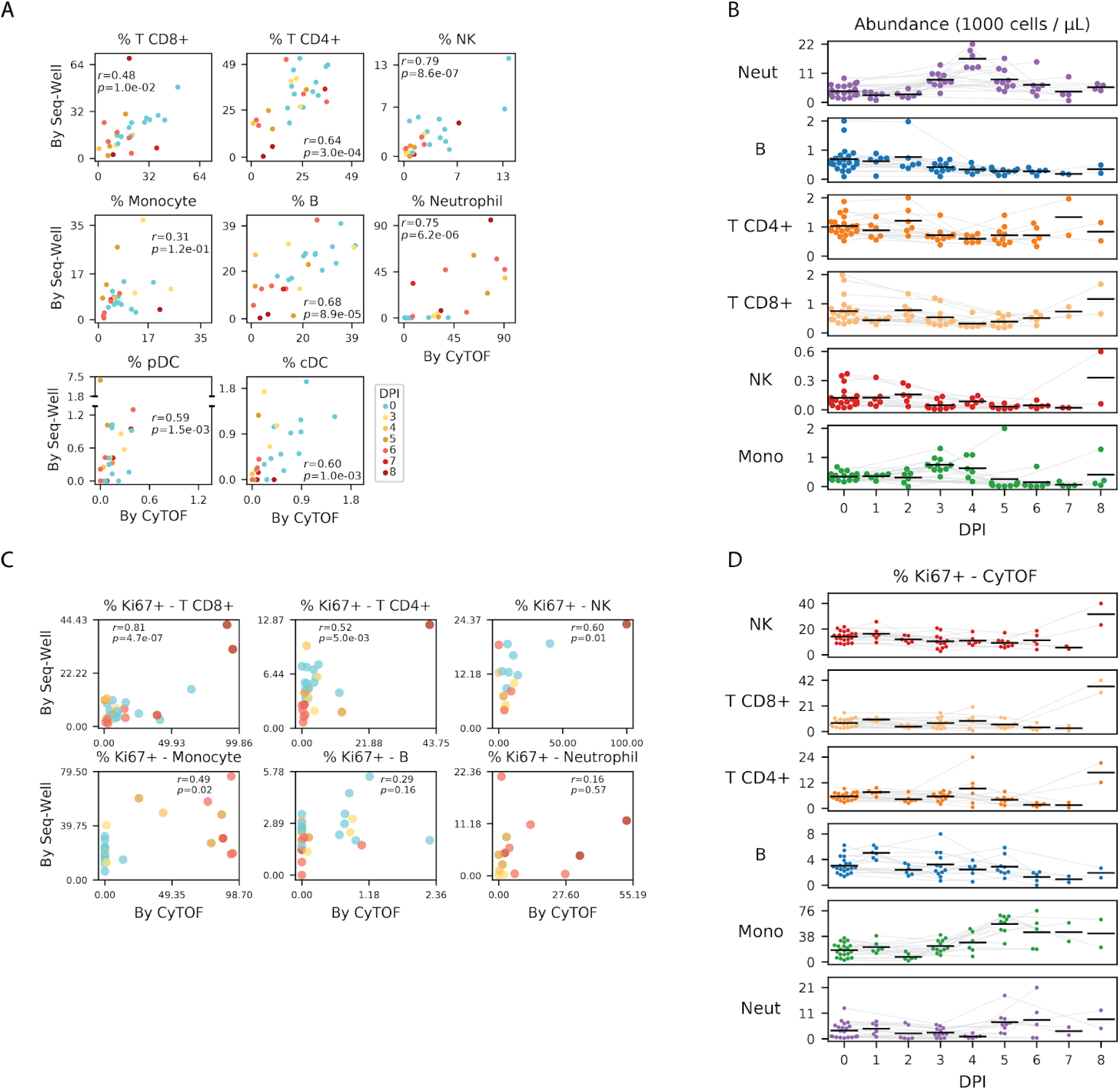
Estimates of cell-type abundance and proliferation over the time course. Related to Figure 2. (**A**) Scatter plot of the percentage of cells of each cell type in a sample, inferred from CyTOF (*x*-axis) or Seq-Well (*y*-axis), for several cell types (panels). Each dot represents a sample colored by DPI. Pearson correlation coefficients (*r*) and *p*-value are provided. (**B**) Estimates of the abundance of each cell type (rows) for each NHP (individual markers) in units of 1000 cells per µL of whole blood, based on integration of CyTOF and complete blood count (CBC) information. The mean value of each DPI is indicated with a black line. Gray lines connect serial samples from the same NHP. (**C**) Scatter plots of the percentage of Ki67-positive cells in a sample inferred from Seq-Well (*x*-axis) or CyTOF (*y*-axis) for several cell types (panels). Each dot represents a sample colored by DPI. Cells with smoothed expression of *MKI67* (the gene coding for Ki67) above 0.1 are called Ki67 positive by Seq-Well. Cells with CyTOF intensity above 1.8 (arbitrary units) are called Ki67 positive by CyTOF. (**D**) Estimates of the percentage of Ki67 positive cells (CyTOF intensity > 1.8) of each cell type (rows) for each animal replicate (markers). The mean value of each DPI is indicated with a black line. Gray lines connect serial samples from the same NHP.

**Figure S6.**
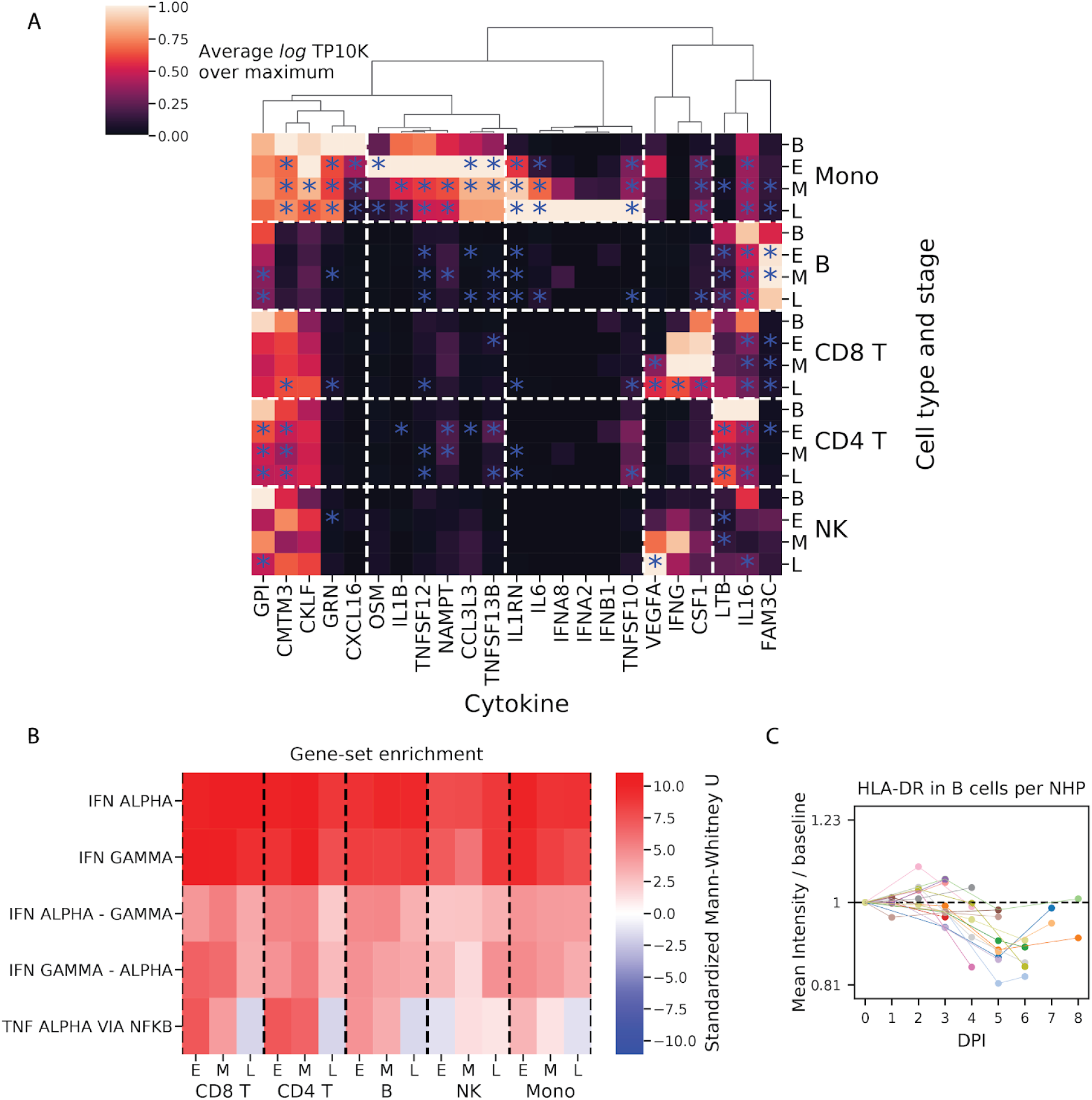
Quantification of cytokine expression and enrichment of response signatures. Related to Figures 3 and 4. (**A**) Average expression values (natural log TP10K) of literature-annotated cytokines (columns) across cell types and stages of acute EVD (rows). Values are plotted as a ratio relative to the maximum across cell types and stages. Values that are statistically different from baseline (*p* < .05) are indicated with a blue star. (**B**) Heatmap of rank-sum test statistics for comparison of differential expression log-fold changes of genes in a gene set (rows) compared to genes not in the gene set. The log fold-changes were defined from differential expression profiles of each cell type at each EVD stage (columns) relative to baseline. Five gene sets were tested -- three from the Hallmark database (IFN ALPHA, IFN GAMMA, and TNF ALPHA VIA NFKB) (Liberzon et al., 2015) and 2 constructed from the hallmark sets, as uniquely IFNα-regulated genes in “IFN ALPHA” but not “IFN GAMMA” (“IFN ALPHA - GAMMA”), and vice versa for uniquely IFNγ-regulated (“IFN GAMMA - ALPHA”). See also **Table S3**. (**C**) Fold change (log_2_ scale) in average HLA-DR CyTOF intensity on B cells at each DPI relative to baseline for each PBMC sample. Colored lines connect serial samples from the same NHP.

**Figure S7.**
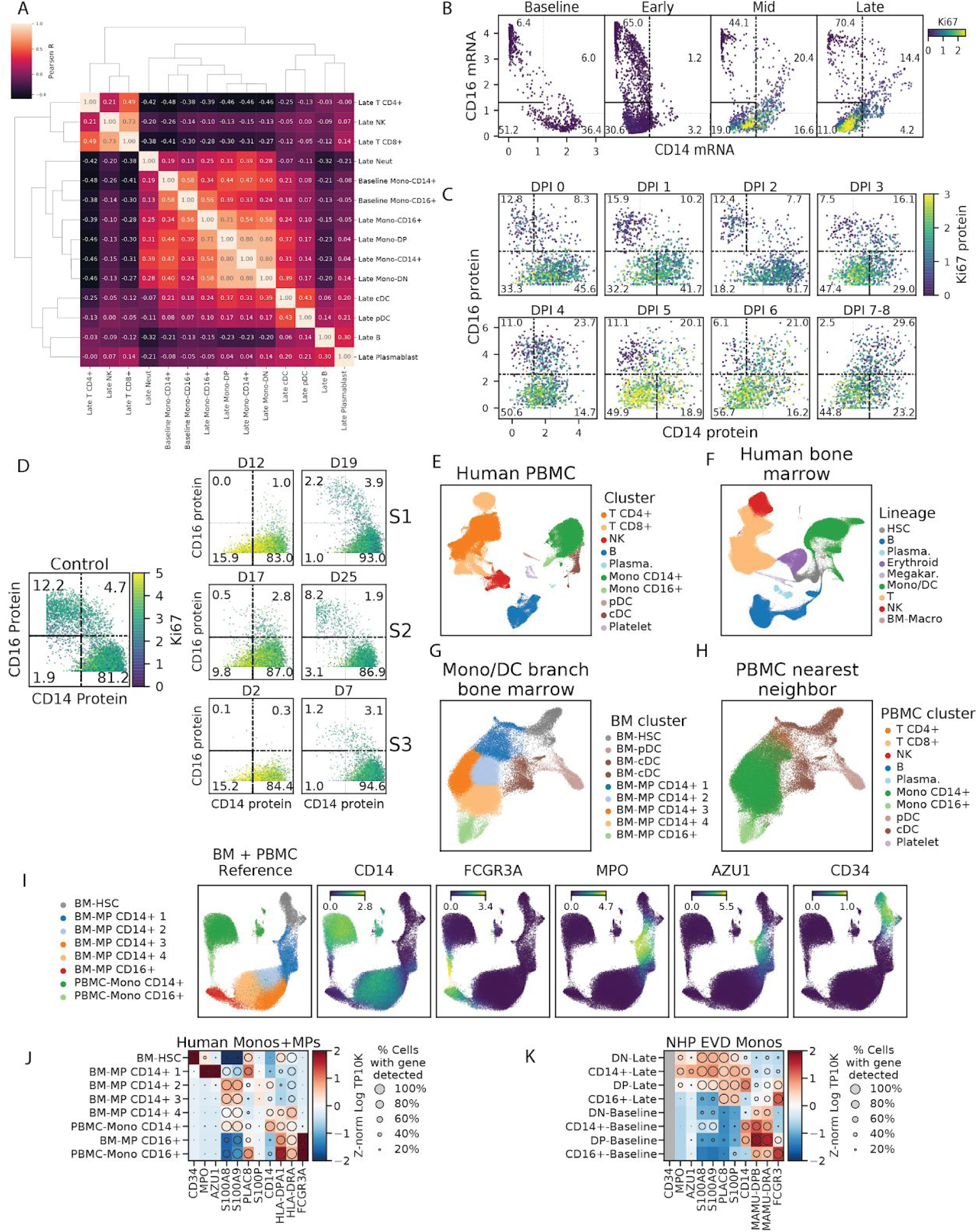
Extended characterization of interferon and double-negative (DN) CD14- CD16- monocytes. Related to Figure 5. (**A**) Clustermap of pairwise Pearson correlations between cell type clusters at baseline and late EVD. Correlations are computed on average log TP10K expression values of overdispersed genes. DN and DP monocytes at late EVD are more similar to monocytes (including baseline CD14+s) than other cell types. (**B**) Scatter plot of MAGIC-smoothed expression values (log TP10K) of *CD14* and *CD16* for monocytes in baseline, early, mid, and late disease stages. Cells are colored by smoothed expression levels of *MKI67* (the gene coding for Ki67 protein). Boxes indicate the CD14+, CD16+, DN, and DP subsets described in the text, and the numbers denote the percentage of cells falling into each subset. (**C**) Scatter plot of protein expression (CyTOF intensity) of CD14 and CD16 for 1000 randomly sampled monocytes at each DPI. Cells are colored by Ki67 expression. Boxes indicate the CD14+, CD16+, DN, and DP subsets described in the text, and the numbers denote the percentage of cells falling into each subset. (**D**) Scatter plot of protein expression (CyTOF intensity) of CD14 and CD16 for monocytes during human EVD. Left: monocytes from healthy human controls. Right: monocytes from 3 EVD cases (S1, S2, and S3) at various days post symptom onset. Cells are colored by Ki67 marker intensity. Boxes indicate the CD14+, CD16+, DN, and DP subsets described in the text and the numbers denote the percentage of cells falling into each subset. (**E**) UMAP embedding of healthy human PBMCs dataset, colored by annotated cluster assignment, based on known marker genes. (Plasma. - Plasmablast). (**F**) UMAP embedding of healthy bone marrow cells, colored by cluster assignment, based on marker genes. (HSC - hematopoietic stem cell, Plasma. - Plasmablast, Megakar. - Megakaryocyte, Mono/DC - monocyte and dendritic cell, BM-Macro - bone marrow resident macrophage). (**G**) UMAP embedding of sub-clustered HSC and monocyte/dendritic lineage cells. (BM - bone marrow, MP - monocyte progenitor) (**H**) Same UMAP embedding as Figure S7G, but cells are colored by the cluster identity of their nearest neighbor in the human PBMC dataset (Figure S7E). (**I**) UMAP embedding of the merged reference dataset of healthy bone marrow HSCs and monocyte lineage cells and PBMCs. Left sub-panel is colored by cluster assignment. Right sub-panels are colored by marker gene expression (natural log TP10K). (**J**) Expression profiles of selected genes for human bone marrow monocyte progenitors (BM-MPs) and human circulating monocytes (PBMC-Monos). Circle area represents the percentage of cells in which the gene was detected, and color denotes the average expression (Z-normalized natural log TP10K). (**K**) Expression profiles of selected genes for NHP monocyte subsets at baseline or late EVD for orthologs of the genes in (J). Circle area represents the percentage of cells in which the gene was detected, and color denotes the average expression level (Z-normalized natural log TP10K). CD34 is grayed out because it is detected in fewer than 10 cells.

**Figure S8.**
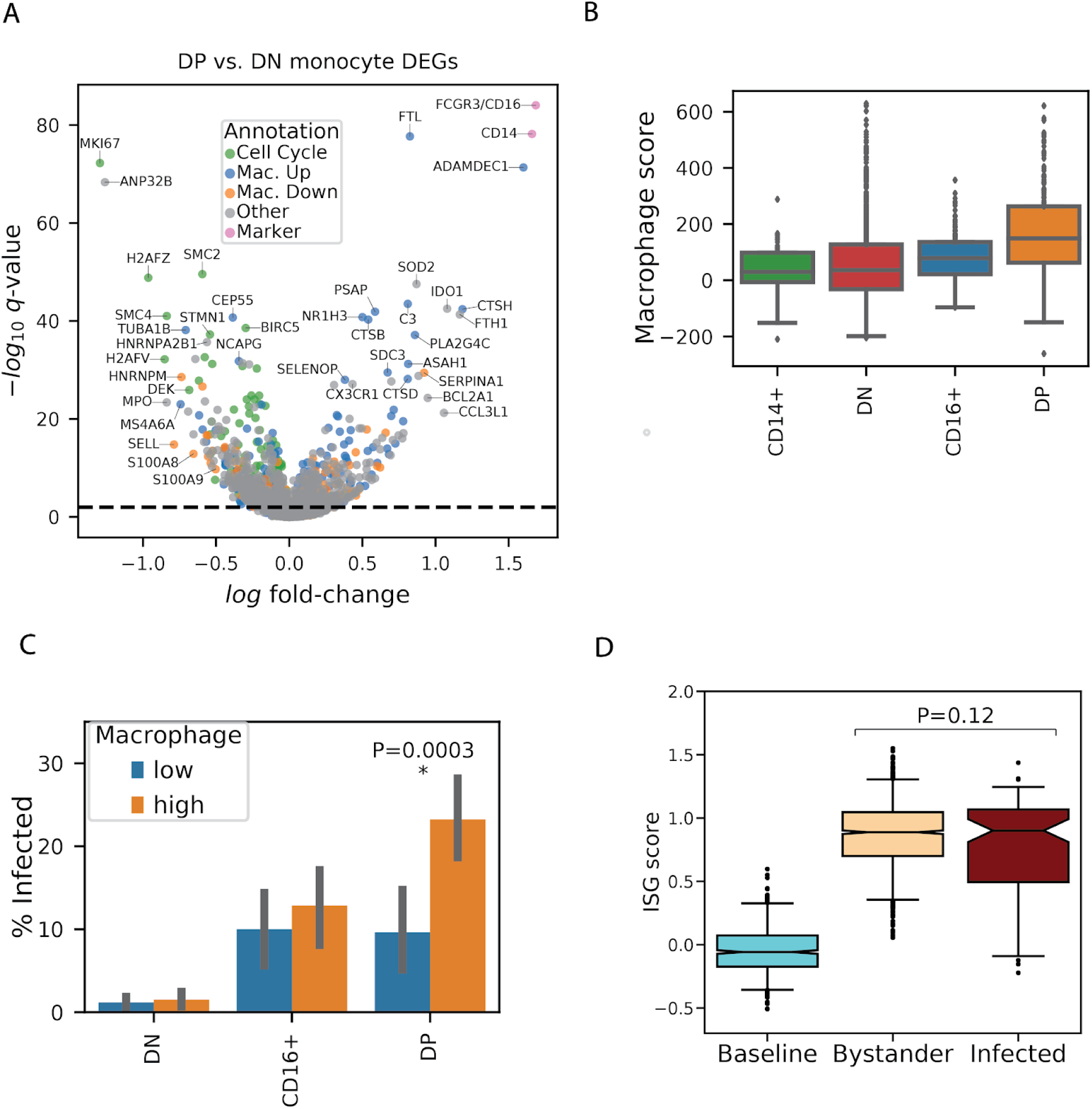
Extended characterization of gene expression signals associated with EBOV infection status in monocytes. Related to Figure 5. (**A**) Volcano plot of differentially expressed genes between double positive and double negative monocyte subsets from DPI 5–8. Genes are colored by membership in cell cycle, macrophage up-regulated (Mac. Up), macrophage down-regulated (Mac. Down), or marker (*CD14*, *CD16*) gene sets. See also **Table S5**. (**B**) Macrophage scores for monocytes in late EVD for each subset. Boxes denote the median and interquartile range, and whiskers denote the 2.5th and 97.5th percentiles. (**C**) Percentage of infected monocytes in each subset in late disease, stratified by low or high macrophage score (below or above the median of monocytes from all subsets). Height of the bar denotes the mean, and error bars denote 95% bootstrap confidence intervals for the mean. There are no infected monocytes in the CD14+ subset. (**D**) ISG scores of monocytes at baseline, and uninfected bystanders or infected cells in late stage EVD (DPI 6–8). Boxes denote the median and interquartile range, and whiskers denote the 2.5th and 97.5th percentiles.

**Figure S9.**
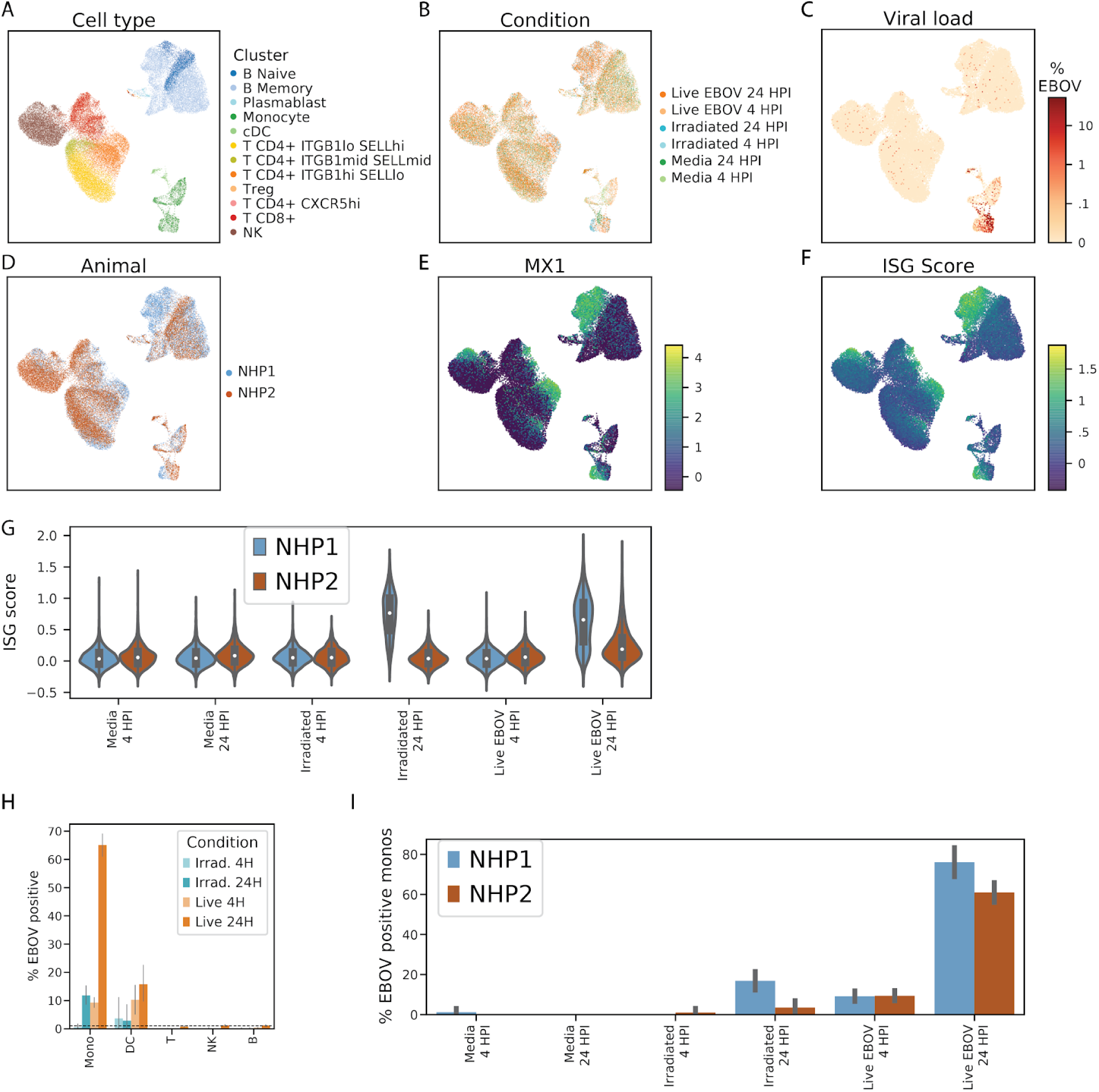
Overview of the *ex vivo* EBOV infection dataset. Related to Figure 6. (**A–F**) Uniform Manifold Approximation and Projection (UMAP) embedding of Seq-Well data colored by annotated cluster assignment (A), treatment condition (B), viral load (C), NHP donor (D), *MX1* gene expression (natural log TP10K) (E), and interferon stimulated gene (ISG) score (F). (**G**) Distributions of ISG scores across monocytes from each treatment condition, stratified by NHP donor. The central white marker denotes the median and the black bar denotes the interquartile range. (**H**) Estimated percentage of infected cells of each cell type in the *ex vivo* dataset. The dashed line denotes the 1% false positive rate threshold used for calling infected cells. (**I**) Percentage of EBOV-positive monocytes from each *ex vivo* treatment condition, stratified by NHP donor. Height of the bar denotes the mean, and error bars denote 95% bootstrap confidence intervals for the mean.

**Figure S10.**
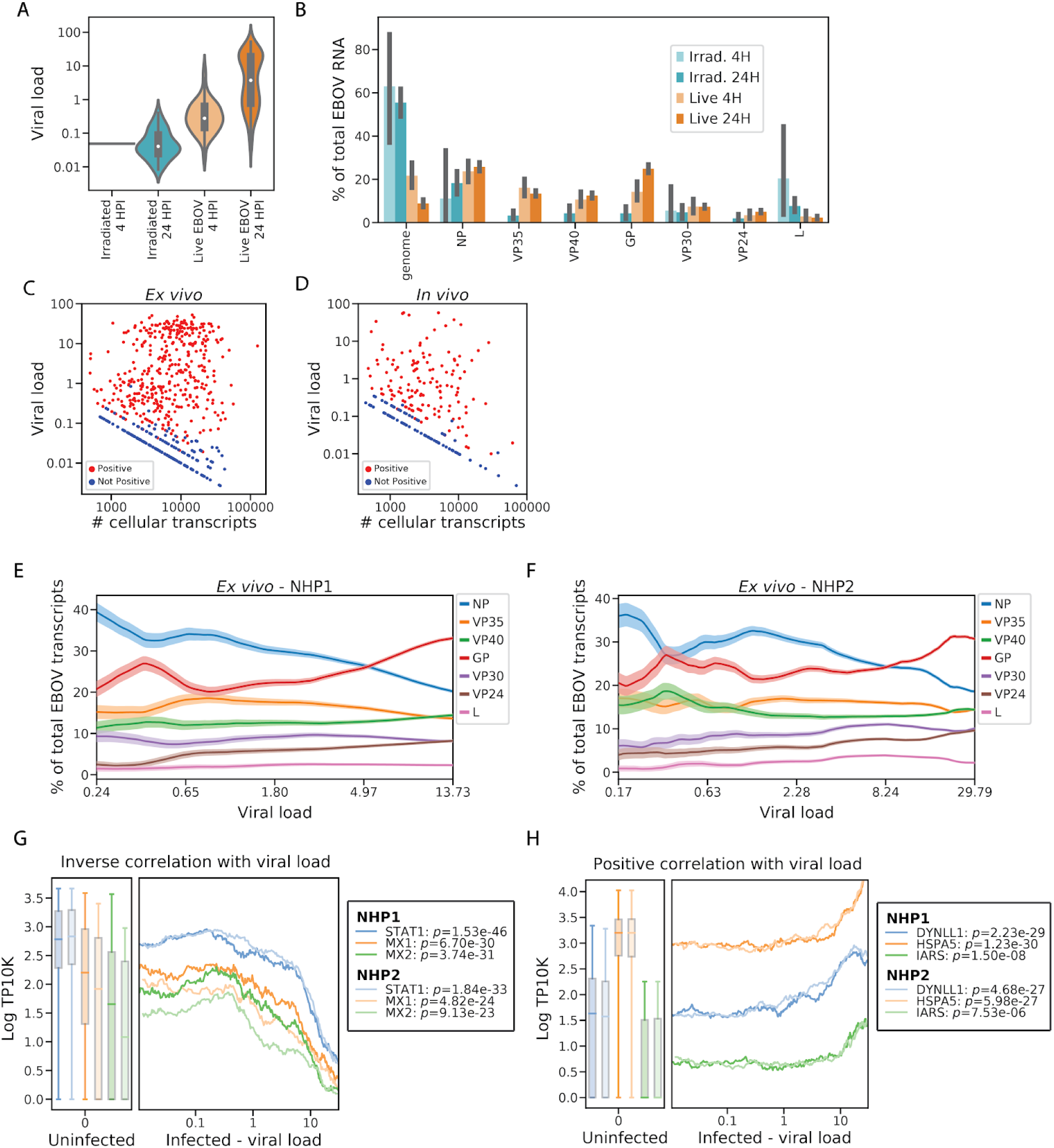
EBOV infection dynamics in the *ex vivo* dataset. Related to Figures 6 and 7. (**A**) Distributions of viral loads across monocytes from different treatment conditions. The central white marker denotes the median and the black bar represents the interquartile range. (**B**) Estimated percentage of EBOV transcripts derived from the EBOV genome or each EBOV gene, out of total viral RNA, stratified by treatment conditions. Prior to averaging, the counts of EBOV genes for each cell was normalized to sum to one, so each cell contributes uniformly to the proportion, regardless of its total number of EBOV transcripts. Height of the bar denotes the mean, and error bars denote 95% bootstrap confidence intervals for the mean. (**C and D**) Scatter plot of total transcripts (unique molecular identifiers) detected in a cell (*x*-axis, log_10_ scale) against viral load (*y*-axis, log_10_ scale) for cells with one or more viral reads *ex vivo* (**C**) or *in vivo* (D). Cells called as infected are colored in red and otherwise colored in blue. (**E and F**) Relative proportion of each EBOV gene versus viral load (log_10_ scale) *ex vivo* for cells from donor NHP1 (E) or NHP2 (F). All monocytes were ordered by viral load and the percentage of each viral gene was averaged over a 50-cell sliding window. Color bands denote the mean ± 1 SD. (**G and H**) Association between gene expression and viral load for selected negatively (G) and positively (G) associated host genes in monocytes, 24 HPI after inoculation with live virus *ex vivo*. In the left sub-plots, distributions of gene expression in uninfected bystander cells are shown as a boxplot with boxes denoting the interquartile range, and whiskers denoting the 2.5th and 97.5th percentiles. In the right sub-plots, we ordered infected cells by viral load (log_10_ scale) and averaged gene expression (natural log TP10K) over a 100-cell sliding window. Curves and box-plots are shown separately for the 2 donor NHPs. *p*-values for the Spearman correlation between viral load and gene expression are listed for each NHP donor in the legend.

## Supplementary Tables

**Table S1. Overview of samples analyzed by CyTOF and Seq-Well. Related to Figure 1.**

Description of each NHP PBMC sample profiled in the study for CyTOF (tab 1) and Seq-Well (tab 2). Includes the animal ID, day post infection, number of cells passing quality control filters, the processing batch, and other technical factors such as whether the sample was processed fresh or underwent freeze-thaw, was treated with DASH, or included an MDCK cell spike-in. Subsequent tabs for the final counts of each broad cell type cluster for CyTOF (tab 3) and Seq-Well (tab 4).

**Table S2. Differential expression profiles of cell types at each EVD stage relative to baseline. Related to Figure 3.**

Log fold-change, *p*-values, and *q*-values for MAST differential expression tests comparing gene expression levels of cells at each EVD stage relative to baseline, for each cell type.

**Table S3. Gene-set enrichment testing of differential expression modules. Related to Figure 3.**

The “geneset_enrichment” tab includes the rank, gene set name, odds ratio (OR), and FDR-corrected q-value for the top 10 gene-sets associated with each differential expression module (ranked by q-value) based on Fisher’s exact tests. The “module_membership” tab includes the module assignment for all genes tested for differential expression including insignificant genes labeled as “Not significant”. The “ISG_set_forScoring” tab includes differentially expressed genes in the “global up” module that were annotated as interferon stimulated genes (ISGs) and subsequently used for computing the ISG score. The Unique_IFN_Alpha_Set species the genes included in the HALLMARK_INTERFERON_ALPHA_RESPONSE gene set but not the HALLMARK_INTERFERON_GAMMA_RESPONSE gene set, and Unique_IFN_Gamma_Set specifies the converse gene set.

**Table S4. Differential expression results of infected vs. bystander monocytes, in vivo, at DPI 5–8. Related to Figure 5.**

Log fold-change, *p*-values, and *q*-values for differential expression tests comparing gene expression levels between infected and bystander monocytes in late EVD. Also includes annotations of whether each gene is associated with macrophage differentiation or interferon response. Subsequent tabs specify the published gene lists associated with in vitro macrophage differentiation that were used for gene set enrichment, as well as the contingency tables and hypothesis tests associated with each enrichment test.

**Table S5. Differential expression between DP and DN monocytes in late EVD. Related to Figure S8.**

Log fold-change, *p*-values, and *q*-values for differential expression tests comparing gene expression levels between DP and DN monocytes in late EVD. Also includes annotations of whether each gene is associated with macrophage differentiation, interferon response, or cell cycle.

**Table S6. Association between host gene expression and viral load for infected monocytes in vivo and ex vivo. Related to Figure 7.**

Log fold-change, *p*-values, and *q*-values for differential expression tests of association between log10 viral load and gene expression for cells containing ≥1 viral read. Tab 1 is for *in vivo* (cells in DPI 5-8), and tab 2 is for *ex vivo* (cells treated with live virus and collected at 24 hours post inoculation). Tabs 3 and 4 show the gene set enrichment results for separate Fisher’s exact tests of genes that are significantly positively or negatively correlated with intracellular viral load (*P* < .05) including enrichment odds ratios (OR), p-values, and q-values.

**Table S7. Antibody panel for cell staining prior to CyTOF. Related to Key Resources Table.**

